# LUZP1, a novel regulator of primary cilia and the actin cytoskeleton, is altered in Townes-Brocks Syndrome

**DOI:** 10.1101/721316

**Authors:** Laura Bozal-Basterra, María Gonzalez-Santamarta, Aitor Bermejo-Arteagabeitia, Carolina Da Fonseca, Olatz Pampliega, Ricardo Andrade, Natalia Martín-Martín, Tess C Branon, Alice Y Ting, Arkaitz Carracedo, Jose A. Rodríguez, Felix Elortza, James D. Sutherland, Rosa Barrio

## Abstract

Primary cilia are sensory organelles that are crucial for cell signaling during development and organ homeostasis. Cilia arise from the centrosome and their formation is governed by numerous regulatory factors. We show that the leucine-zipper protein LUZP1 localizes to the pericentriolar material and actin cytoskeleton. Using TurboID proximity labeling and pulldowns, LUZP1 associates with factors linked to centrosome and actin filaments. Loss of LUZP1 reduces F-actin levels, facilitating ciliogenesis and altering Sonic Hedgehog signaling, pointing to a key role in the cytoskeleton-cilia interdependency. Moreover, we show that LUZP1 interacts with a truncated form of the transcription factor SALL1 that causes Townes-Brocks Syndrome. TBS is characterized by digit, heart and kidney malformations and is linked in part to defective cilia. Truncated SALL1 increases the ubiquitin proteasome-mediated degradation of LUZP1. Alteration of LUZP1 levels may be a contributing factor to TBS, suggesting possible therapies using modulators of cilia and cytoskeletal function.

## INTRODUCTION

Primary cilia are sensory organelles that have a crucial role in cell signaling, polarity and protein trafficking during development and organ homeostasis. Importantly, the involvement of primary cilia in the above-mentioned processes is frequently due to its role in Sonic Hedgehog (Shh) pathway regulation (1). Briefly, Shh activation through its receptor PTCH1 leads to ciliary enrichment of the transmembrane protein Smoothened (SMO), with concomitant conversion of the transcription factor GLI3 from a cleaved repressor form to a full-length activator form, leading to activation of Shh target genes. Two such genes are *PTCH1* and *GLI1* (encoding the Shh receptor and a transcriptional activator, respectively), exemplifying the feedback and fine-tuning of the Shh pathway.

Cilia arise from the centrosome, a cellular organelle composed of two barrel-shaped microtubule-based structures called the centrioles. Primary cilia formation is very dynamic throughout the cell cycle. Cilia are nucleated from the mother centriole (MC) at the membrane-anchored basal body upon entry into the G0 phase, and they reabsorb as cells progress from G1 to S phase, completely disassembling in mitosis (2). Centrioles are surrounded by protein-based matrix pericentriolar material (PCM) (3, 4). In eukaryotic cells, PCM proteins are concentrically arranged around a centriole in a highly organized manner (5–8). Based on this observation, proper positioning and organization of PCM proteins may be important for promoting different cellular processes in a spatially regulated way (9). Not surprisingly, aberrations in the function of PCM scaffolds are also associated with many human diseases, including cancer and ciliopathies (10, 11).

Cilia assembly and disassembly are regulated by diverse factors, including the main cilia suppressor proteins CCP110 and CEP97 and the actin cytoskeleton. CCP110 and CEP97 form a complex that, when removed from the MC, allows ciliogenesis (12). The regulation of actin dynamics is also considered a major ciliogenesis driver in cycling cells (13).

Ciliary dysfunction often results in early developmental problems including hydrocephalus, neural tube closure defects (NTD) and left-right anomalies (14). These features are often reported in a variety of diseases, collectively known as ciliopathies, caused by failure of cilia formation and/or cilia-dependent signaling (15). In the adult, depending on the underlying mutation, ciliopathies present a broad spectrum of phenotypes comprising cystic kidneys, polydactyly, obesity or heart malformation.

Townes-Brocks Syndrome (TBS1 [MIM: 107480]) is an autosomal dominant genetic disease caused by mutations in *SALL1*, characterized by the presence of imperforate anus, dysplastic ears, thumb malformations, and often with renal and heart impairment, among other symptoms (16, 17), features seen in the ciliopathic spectrum. It has been recently demonstrated that primary cilia defects are contributing factors to TBS aetiology (18). Truncated SALL1, either by itself or in complex with the SALL1 full length form (SALL1^FL^), can interact with CCP110 and CEP97. As a consequence, those negative regulators disappear from the MC and ciliogenesis is promoted (18). Truncated SALL1 likely interferes with multiple factors to give rise to TBS phenotypes. Here we focus on LUZP1, a leucine-zipper motif containing protein that was identified by proximity proteomics as an interactor of truncated SALL1 (18).

LUZP1 has also been identified as an interactor of ACTR2 (ARP2 actin related protein 2 homologue) and filamin A (FLNA) and, recently, as an actin cross-linking protein (19, 20). Furthermore, LUZP1 shows homology to FILIP1, a protein interactor of FLNA and actin (21, 22). Interestingly, mutations in *Luzp1* resulted in cardiovascular defects and cranial NTD in mice (23), phenotypes within the spectrum of those seen in TBS individuals and mouse models of dysfunctional cilia, respectively (16, 17, 24–27). LUZP1 was found to be mainly localized to the nuclei of brain neurons in mice and to have a crucial role in embryonic brain development (23, 28, 29). Both the planar cell polarity/Wingless-Integrated (Wnt) pathway and the Sonic Hedgehog (Shh) pathway are influenced by the presence of functional cilia and regulate neural tube closure and patterning (30–32). Remarkably, ectopic SHH was observed in the dorsal lateral neuroepithelium of the *Luzp1****^−/−^*** mice (23). However, in spite of the phenotypic overlaps, a link between LUZP1 and ciliogenesis had not been previously investigated.

Here we demonstrate that LUZP1 is associated with centrosomal and actin cytoskeleton-related proteins. We also demonstrate that LUZP1 localizes to the PCM, actin cytoskeleton and the midbody, providing evidence towards its regulatory role on actin dynamics and its subsequent impact on ciliogenesis. Notably, we demonstrate that Luzp1^−/−^ cells exhibit reduced polymerized actin, longer primary cilia, higher rates of ciliogenesis and increased Shh signaling. Furthermore, TBS-derived primary fibroblasts show a reduction in LUZP1 and actin filaments (F-actin), possibly through SALL1-regulated LUZP1 degradation via the ubiquitin (Ub)-proteasome system (UPS). Altogether, these results indicate that LUZP1 participates in ciliogenesis and maintenance of the actin cytoskeleton and might contribute to the aberrant cilia phenotype in TBS.

## RESULTS

### SALL1 interacts with LUZP1

We have previously shown that a truncated and mislocalized form of SALL1 present in TBS individuals (SALL1^275^) can interact aberrantly with cytoplasmic proteins. (18). LUZP1 was found among the most enriched proteins in the SALL1^275^ interactome. We confirmed this finding by independent BioID experiments analyzed by Western blot using a LUZP1-specific antibody (Figure 1A and Figure 1-figure supplement 1). To further characterize the interaction of LUZP1 with SALL1, we performed pulldowns with tagged SALL1^275^-YFP in HEK 293FT cells. Our results showed that endogenous LUZP1 was able to interact with SALL1^275^, confirming our proximity proteomics data (Figure 1B, lane 6, and Figure 1-figure supplement 1). The interaction with SALL1^275^ persisted in presence of overexpressed *SALL1^FL^* (Figure 1B, lane 9, and Figure 1-figure supplement 1), suggesting that the possible heterodimerization of the truncated and FL forms does not inhibit the interaction with LUZP1. Of interest, LUZP1 also interacts with SALL1^FL^ when overexpressed alone (Figure 1B, lane 7 and Figure 1-figure supplement 1). These results support the notion that the truncated form of SALL1 expressed in TBS individuals, either by itself or in complex with the FL form, can interact with LUZP1.

**Figure 1.**
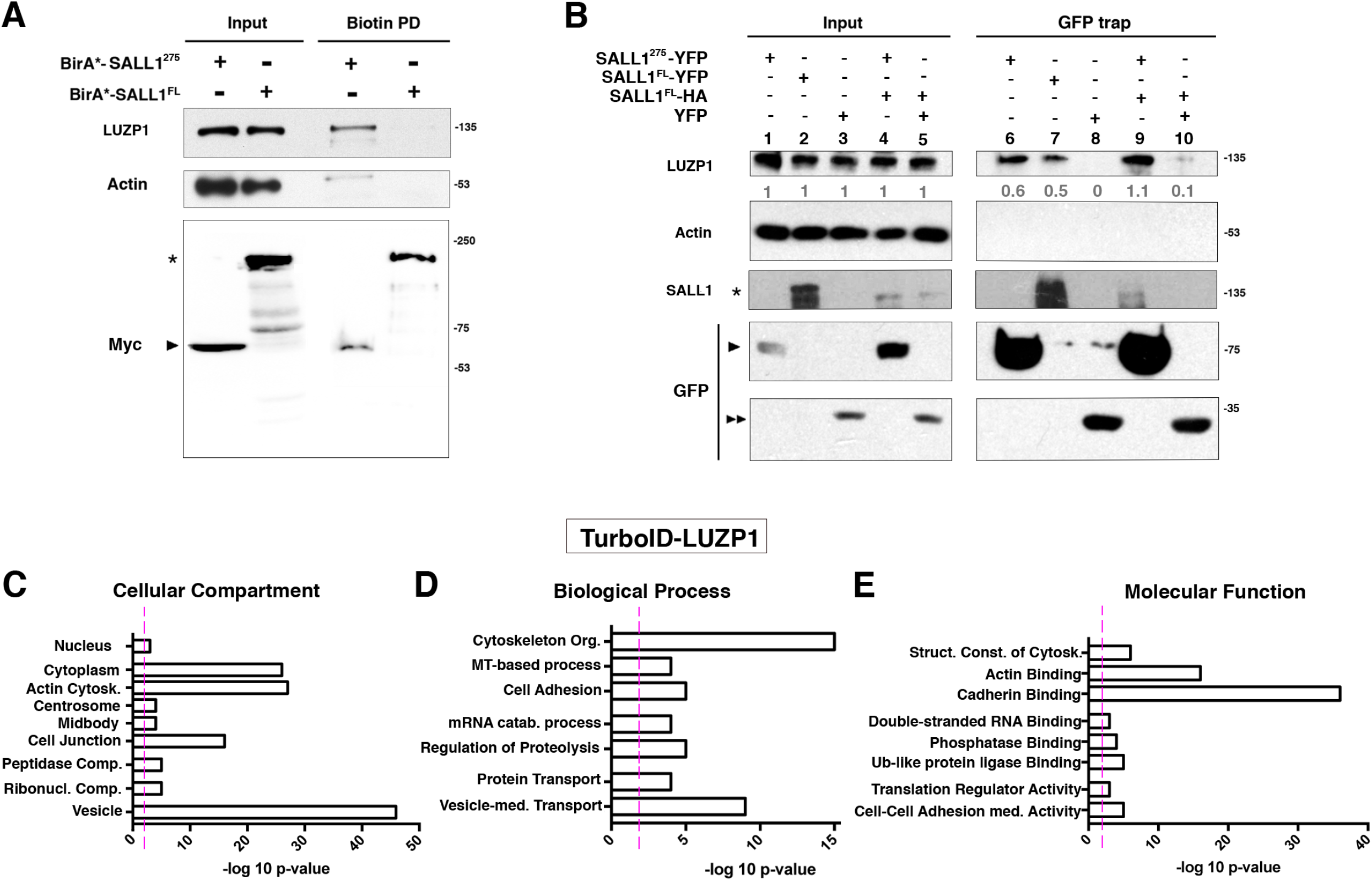
Proximity proteomics reveals LUZP1 interacting with truncated SALL1 and with centrosome and actin cytoskeleton-related proteins. (**A**) Western blot analysis of BioID, biotin pulldown (PD) of HEK 293FT cells transfected with Myc-tagged BirA*-SALL1^275^ (BirA*-SALL1^275^) or BirA*-SALL1^FL^. Specific antibodies (LUZP1, actin, Myc) were used as indicated. Anti-Myc antibody detected the self-biotinylated form of BirA*-SALL1^FL^ (asterisk) or BirA*-SALL1^275^ (black arrowhead). (**B**) Western blot of inputs or GFP-Trap pulldowns performed in HEK 293FT cells transfected with *SALL1^275^-YFP* (lanes 1 and 6), *SALL1^FL^-YFP* (lanes 2 and 7), *YFP* alone (lanes 3 and 8), *SALL1^275^-YFP* together with *SALL1^FL^-2xHA* (SALL1^FL^-HA; lanes 4 and 9) or SALL1^FL^-HA together with YFP alone (lanes 5 and 10). Specific antibodies (LUZP1, actin, SALL1) were used as indicated. Numbers under LUZP1 panel result from dividing band intensities of each pulldown by their respective input levels. One asterisk indicates BirA*-SALL1^FL^ or SALL1^FL^-YFP, one black arrowhead SALL1^275^-YFP and two black arrowheads YFP alone. Molecular weight markers (kDa) are shown to the right. Actin was used as loading control. Blots shown are representative of three independent experiments. (**C-E**) Graphical representation of the -log_10_ of the P-value for each of the represented GO terms of the TurboID performed on hTERT-RPE1 stably expressing near endogenous levels of *FLAG-TurboID-LUZP1*: Cellular Compartment (C), Biological Process (D) and Molecular Function (E). Cytosk.: cytoskeleton; Comp.: complex; Ribonucl.: ribonucleoprotein; Org.: organization; MT: microtubules; catab.: catabolism; med.: mediated; Struct. Const. of Cytosk: structural constituent of cytoskeleton; Ub: ubiquitin. Pink dotted line represents the cutoff of P value <0.01. The following figure supplement is available for Figure 1: Figure 1-figure supplement 1. Western blot full pictures for Figure 1.

### LUZP1 proximal interactors enriched for centrosomal and actin cytoskeleton components

To gain some clues into the function of LUZP1, we sought to identify its proximal interactome using the TurboID approach (33). We used hTERT-RPE1 cells stably expressing low levels of FLAG-TurboID-LUZP1, and after a brief biotin-labeling, biotinylated proteins were captured for analysis by liquid chromatography tandem mass spectrometry (LC-MS/MS). 311 high-confidence proximity LUZP1 interactors were identified in at least two replicates (Table S1). With the purpose of obtaining a functional overview of the main pathways associated to LUZP1, a comparative Gene Ontology (GO) analysis was performed with all the hits (Figure 1C-E and Table S1). In the Cellular Component domain, “cytoplasm”, “actin cytoskeleton”, “centrosome”, “midbody”, “cell junction” and “vesicle” terms were highlighted (Figure 1C and Table S1). In the category of Biological Process, LUZP1 proteome shows enrichment in the “cytoskeleton organization”, vesicle-mediated transport and cell adhesion categories among others (Figure 1D and Table S1). With respect to Molecular Function, LUZP1 also showed enrichment in cytoskeleton-related proteins (“structural component of cytoskeleton” and “actin binding” terms; Figure 1E and Table S1). 64 or 138 of the verified or potential, respectively, centrosome/cilia gene products previously identified by proteomic analyzes (34, 35) were found as LUZP1 proximal interactors, supporting the enrichment of centrosome-related proteins among the potential interactors of LUZP1. In addition, 48 of LUZP1 proximal interactors were present among the actin-localized proteins identified by the Human Protein Atlas project based on actin filaments subcellular localization (36).

### LUZP1 localizes to the PCM, with altered levels and distribution in TBS fibroblasts

Based on the interaction of LUZP1 with centrosomal proteins, we examined the subcellular localization of LUZP1 at the centrosome. Immunostainings showed that LUZP1 appeared as a basket-like 3D structure surrounding both centrioles, which were labelled by centrin 2 (CETN2) staining in human RPE1 cells (Figure 2A) as well as in human dermal fibroblasts (Figure 2B and Figure 2-supplementary video 1) and U2OS cells (Figure 2C). Next, to compare LUZP1 with additional centriolar markers, we labelled U2OS cells expressing *YFP-LUZP1* with the distal centriolar markers CCP110 and ODF2 (outer dense fiber of sperm tails 2). We did not observe colocalization of LUZP1 with these markers, indicating that LUZP1 is likely found at the proximal end of both centrioles (Figure 2D). We further imaged LUZP1 along with PCM1 and centrobin, markers of PCM and of the MC, respectively. Interestingly, we observed LUZP1 being surrounded by PCM1 (Figure 2E), while LUZP1 surrounded centrobin at the MC (Figure 2G). The profile histograms confirm that LUZP1 localizes between PCM1 and centrobin (Figure 2F and 2H, respectively), suggesting that LUZP1 might be a novel PCM associated-protein, forming a basket around the proximal end of both centrioles. We also examined LUZP1 localization in the centrosome in synchronized human RPE1-cells. LUZP1 was reduced at the centrosome during G2/M and G0 phases (Figure 2-figure supplement 2). LUZP1 levels increased upon treatment with the proteasome inhibitor MG132 in G0 phase arrested-RPE1 cells.

**Figure 2.**
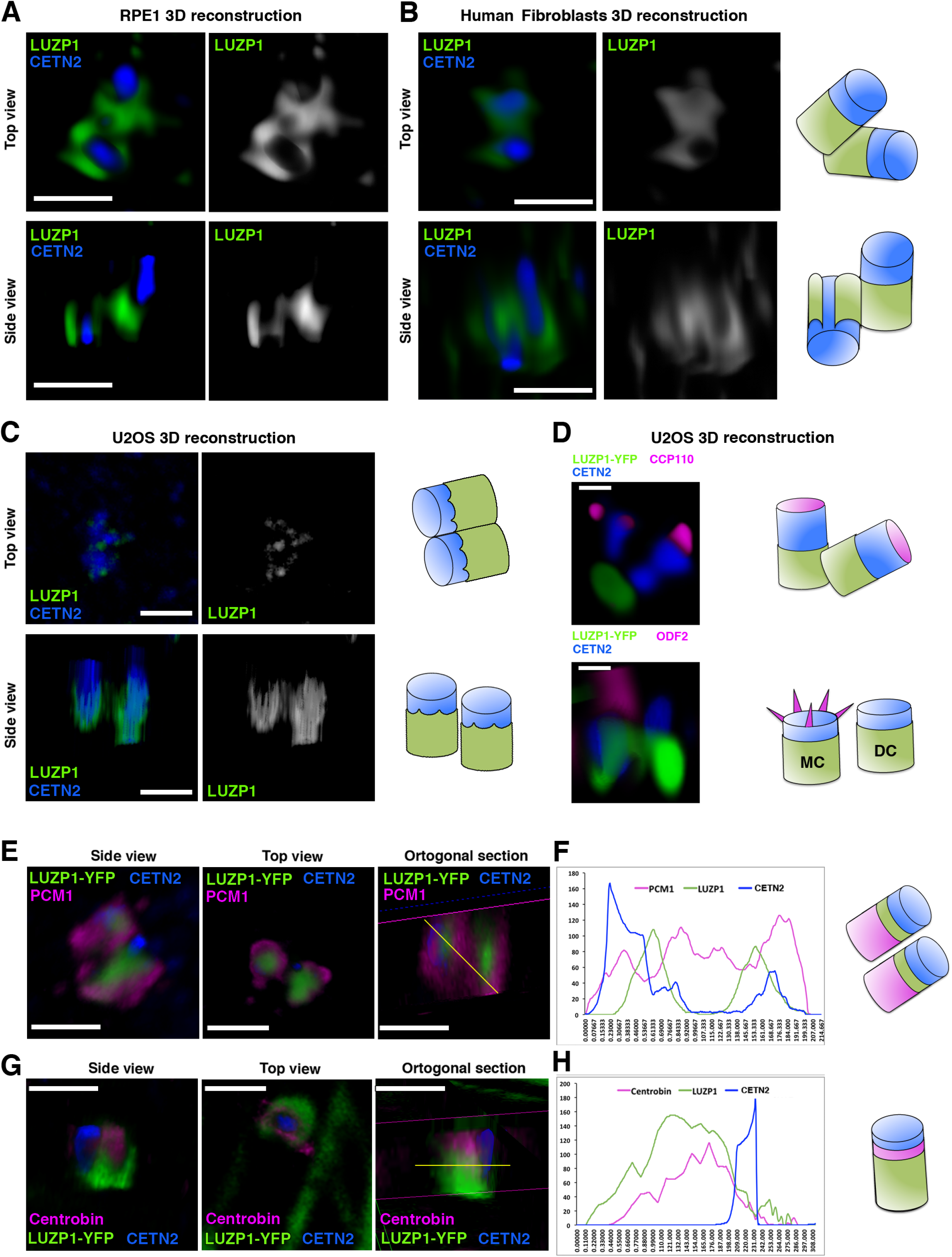
LUZP1 localizes to the proximal end of both centrioles. (**A, B, C**) 2D images of a 3D reconstruction of immunofluorescence micrographs of LUZP1 (green) and Centrin-2 (CETN2, blue) in RPE1 cells (**A**), control fibroblasts (ESCTRL#2) (**B**) or U2OS cells (**C**). (**D**) 2D sections of U2OS cells overexpressing GFP-LUZP1 (green) stained with antibodies against CCP110 (upper panel, magenta) or ODF2 (lower panel, magenta) and CETN2 (blue). (**E, G**) 3D immunofluorescence micrographs of U2OS cells overexpressing LUZP1-YFP (green) stained with antibodies against PCM1 (**E**) or centrobin (**G**) in magenta and CETN2 (blue). Purple lines indicate the orthogonal cuts of the confocal z-stacks sections; yellow lines indicate the quantification point in (**F**) and (**H**). (**F, H**) Plot profile of the orthogonal section in (**E**) or (**G**) showing LUZP1-YFP (green), PCM1 or centrobin (magenta) and CETN2 (blue) intensities along the yellow lines in **E** and **G**, from left to right. Schematic representation of LUZP1 localization at the centrosome was modelled according to their respective micrographs in (**A-G**). Scale bar, 1 µm (**A-D**) or 0.5 µm (**D, E, G**). Imaging was performed using confocal microscopy (Leica SP8, 63x objective). Lightning software (Leica) was applied. The following figure supplement is available for Figure 2: Figure 2-figure supplement 1. Centrosomal localization of LUZP1 along the cell cycle. Figure 2-Supplementary video 1. LUZP1 localization in the centrosome.

To see whether LUZP1 localization and levels are affected in TBS, we checked its subcellular localization using super-resolution microscopy in fibroblasts derived from a TBS individual (TBS^275^; see Materials and Methods) as well as non-TBS controls. Our results showed that LUZP1 was markedly decreased in TBS^275^ cells compared to control cells in non-starved conditions (Figure 3A,B) and that LUZP1 was visualized as two rings that circled each of the centrioles, stained with gamma tubulin, both in control and TBS^275^ cells at the base of primary cilia. We also found LUZP1 localized in scattered dots along the ciliary shaft in starved cells (Figure 3A,B, yellow arrows).

**Figure 3.**
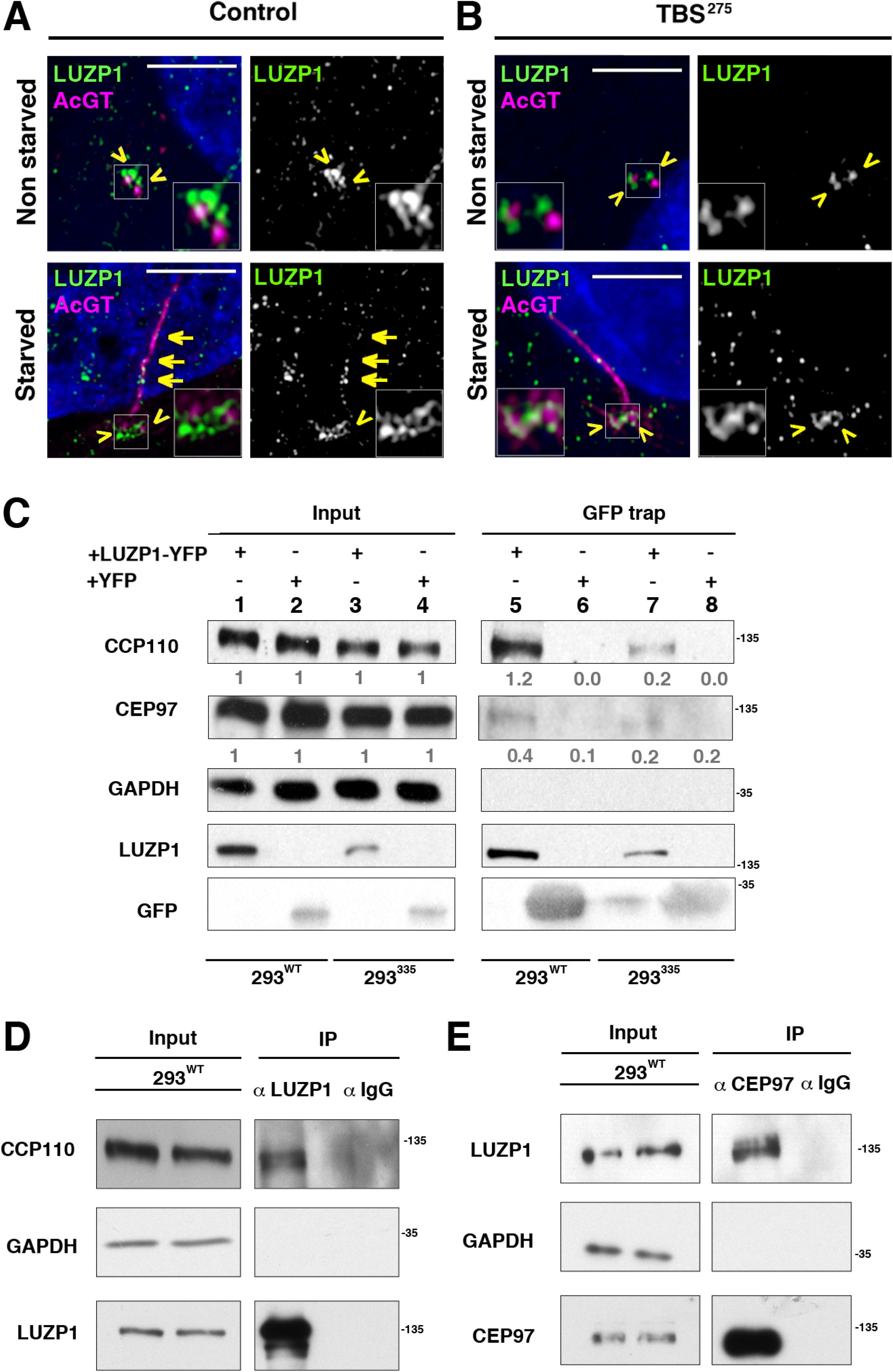
TBS cells show reduction in LUZP1 levels at the centrosome. (**A, B**) Immunofluorescence micrographs of non-starved and starved human-derived control ESCTRL#2 (**A**) and TBS^275^ fibroblasts (**B**) stained with antibodies against endogenous LUZP1 (green, yellow arrows and arrowheads) and acetylated alpha-tubulin and gamma tubulin to simultaneously label the cilia and centrosomes, respectively (magenta). Black and white images show the isolated green channel. Note the reduction of LUZP1 in starved cells and in TBS^275^ compared to control ESCTRL2 fibroblasts. Imaging was performed using confocal super-resolution microscopy (Zeiss LSM 880 Fast Airyscan, 63x objective). AcGT: acetylated and gamma tubulin. Scale bar, 4 µm. (C) Western blot of inputs (lines 1 to 4) and GFP-Trap pulldowns (lines 5 to 8) performed in WT HEK 293FT cells or in 293^335^ *SALL1* mutant cells transfected with *LUZP1-YFP* (lanes 1, 3, 5 and 7) or *YFP* alone (lanes 2, 4, 6 and 8). Numbers under CCP110 and CEP97 panels result from dividing band intensities of each pulldown by their respective input levels. GAPDH was used as loading control. (**D, E**) Co-immunoprecipitation experiments show LUZP1-CCP110 (**D**) and CEP97-LUZP1 (**E**) interactions. Rabbit IgGs were used as immunoprecipitation controls. GAPDH was used as loading and specificity control. In all panels, specific antibodies (LUZP1, GAPDH, CCP110, CEP97, GFP) were used as indicated. Blots shown here are representative of three independent experiments. Molecular weight markers are shown to the right. The following figure supplement is available for Figure 3: Figure 3-figure supplement 1. Western blot full pictures for Figure 3.

### LUZP1 interacts with centrosomal regulators

LUZP1 localization in human cells and proximity labelling experiments suggested that this protein might associate with centrosome-related proteins. We previously found that SALL1^275^-YFP interacted with the centrosome-associated ciliogenesis suppressors, CCP110 and CEP97 (18), so we checked whether LUZP1 may also interact with these factors. Indeed, LUZP1-YFP interacts with CCP110 and CEP97 in both WT HEK 293FT (293^WT^) and in a TBS model cell line, 293^335^ (Figure 3C, lanes 5 and 7, respectively, and Figure 3-figure supplement 1) (18). Less CCP110 and CEP97 was recovered in LUZP1-YFP pulldowns from 293^335^ cells, but this is likely due to the reduced LUZP1-YFP seen in those cells (Figure 3C, Input, lanes 1 and 2 *vs* lane 3 and 4). Beyond pulldowns, we found that immunoprecipitation of endogenous LUZP1 led to co-purification of endogenous CCP110 (Figure 3D and Figure 3-figure supplement 1) and that anti-CEP97 antibodies immunoprecipitated endogenous LUZP1 (Figure 3E and Figure 3-figure supplement 1). These results confirm that, in agreement with the localization of the protein to the centrosome, LUZP1 associates with core centrosomal components.

### LUZP1 localizes to actin and is altered in TBS fibroblasts

In addition to the localization at the centrosome/basal body, LUZP1 also localized to the actin fibers and in the midbody in dividing cells (Figure 4A and Figure 4-figure supplement 2 and 3). We analyzed LUZP1 levels in synchronized human RPE1-cells. Similarly to the changes observed at the centrosome, LUZP1 levels were reduced during the G2/M and GO phases (Figure 4-figure supplement 2). Proteasome inhibition during G0 led to increased LUZP1, suggesting that active degradation occurs in G0 arrested-RPE1 cells. Intriguingly, when LUZP1 levels were examined in TBS^275^ cells, a reduction in both actin-associated LUZP1 and phalloidin-labelled stress fibers was observed when compared to control cells (Figures 4A-C). These results indicate that actin cytoskeleton might be altered in TBS cells. By pulldown assays, we confirmed that LUZP1-YFP interacts with both actin and FLNA (Figure 4D and Figure 4-figure supplement 1). Of note, actin, FLNA and other stress fibers-associated proteins were also found to be associated with LUZP1 by proximity labeling and mass spectrometry (Table S1). To examine whether LUZP1 levels change upon F-actin perturbation, HEK 293FT cells were treated with cytochalasin D (CytD), an actin-polymerization inhibitor. No changes in LUZP1 levels upon actin depolymerisation were observed when cells were lysed in strong lysis conditions (WB5). However, we observed a consistent increase in LUZP1 levels using mild lysis condition in extraction buffer containing 1% Triton X-100 (Figure 4E,F and Figure 4-figure supplement 1). These results reflect that the integrity of the actin cytoskeleton may influence the solubility but not the stability of LUZP1.

**Figure 4.**
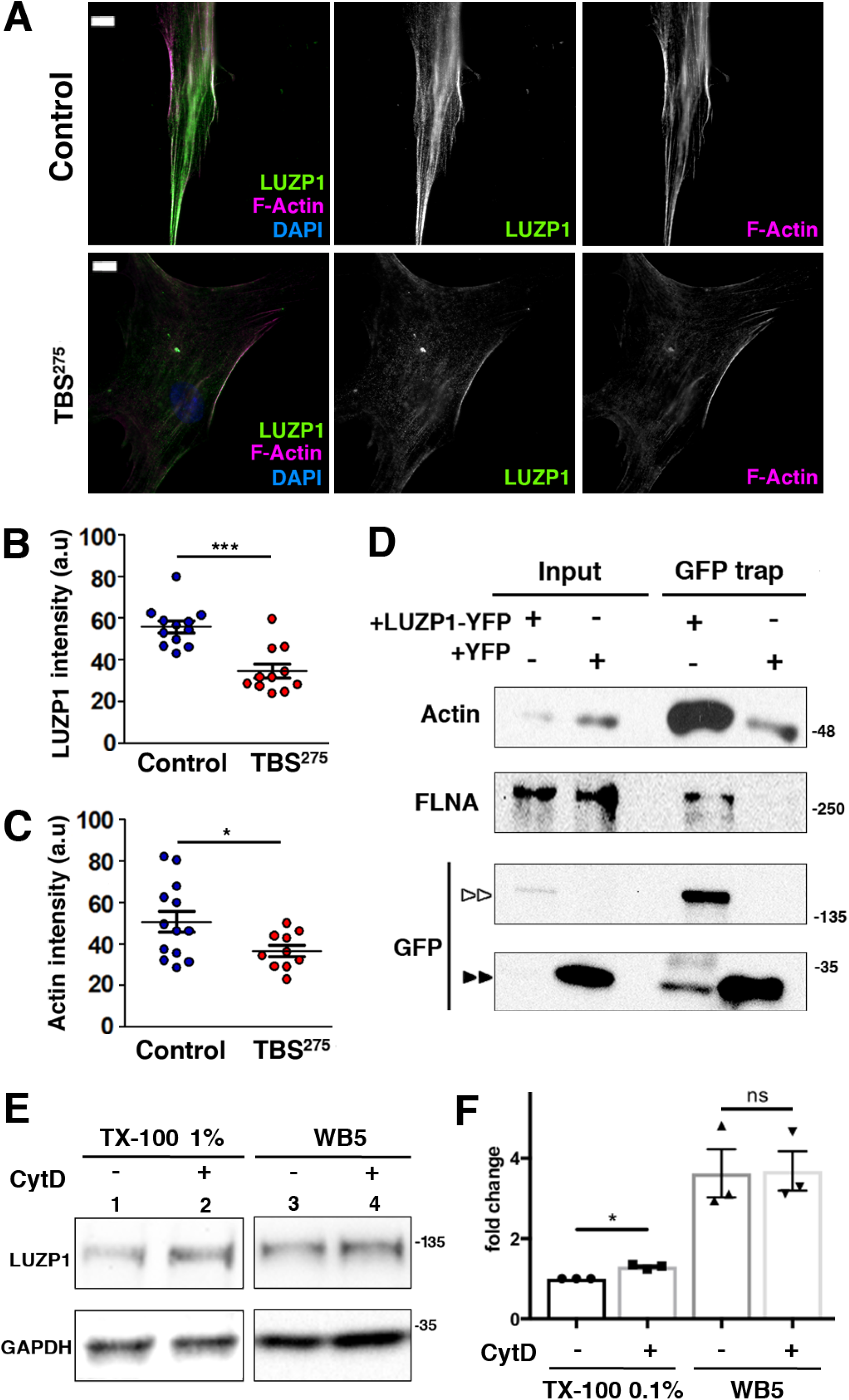
Reduction in LUZP1 is linked to F-actin decrease. (**A**) Immunofluorescence micrographs of control ESCTRL#2 and TBS^275^ human fibroblasts stained with an antibody against endogenous LUZP1 (green), phalloidin to label F-actin (magenta), and counterstained with DAPI to label the nuclei (blue). Black and white images show the single green and magenta channels. Note the overall reduction in LUZP1 and F-actin levels in TBS^275^ compared to control ESCTRL2 fibroblasts. Scale bar, 10 µm. Imaging was performed using widefield fluorescence microscopy (Zeiss Axioimager D1, 63x objective). (**B, C**) Graphical representation of the LUZP1 (**B**) and F-actin (**C**) mean intensities, corresponding to the experiments shown in (**A**); n≥6 micrographs. Three independent experiments were pooled together. P-values were calculated using the unpaired two-tailed Student’s test or U-Mann-Whitney test. (**D**) Western blot of inputs or GFP-Trap pulldowns performed in HEK 293FT cells transfected with *LUZP1-YFP* or *YFP* alone. Anti-GFP antibody detected YFP alone (two black arrowheads) and LUZP1-YFP (two white arrowheads). Blots shown here are representative of three independent experiments. Molecular weight markers are shown to the right. Specific antibodies (LUZP1, GAPDH, CCP110, CEP97, GFP) were used as indicated. (**E**) Western blot of total cell lysates of HEK 293FT treated or not with Cytochalasin D (CytD) in a mild lysis buffer (TX-100 1%, lanes 1, 2) or a strong lysis buffer (WB5, lanes 3, 4). Note the increase in LUZP1 levels upon actin polymerization blockage with CytD, exclusively when cells were lysed on 1% TX-100-based lysis buffer. GAPDH was used as loading control. In (**D**) and (**E**) panels, specific antibodies (LUZP1, GAPDH, actin, FLNA, GFP) were used as indicated. (**F**) Graphical representation of LUZP1 *vs* GAPDH band intensities in (**E**) normalized to lane 1. Graphs represent Mean and SEM of three independent experiments. P-value was calculated using two tailed unpaired Student’s t-test. The following figure supplement is available for Figure 4: Figure 4-figure supplement 1. Western blot full pictures for Figure 4. Figure 4-figure supplement 2. LUZP1 localization at the cytoskeleton along the cell cycle. Figure 4-figure supplement 3. LUZP1 localization at the midbody.

### LUZP1 plays a role in primary cilia formation and F-actin stabilization

Based on the LUZP1 localization at the centrosome, its interaction with centrosomal proteins and the defects in ciliogenesis previously observed in TBS cells (18), we hypothesised that LUZP1 might have a role in cilia formation. To examine this, we analyze ciliogenesis in Shh-LIGHT2 cells, a cell line derived from immortalized mouse NIH3T3 fibroblasts that display primary cilia and carry a Shh luciferase reporter (herein considered as WT fibroblasts) (37). Additionally, using CRISPR/Cas9 gene editing directed to exon 1 of murine *Luzp1*, we generated Shh-LIGHT2 mouse embryonic fibroblasts null for *Luzp1* (Luzp1^−/−^ cells), and for genetic rescue experiments, LUZP1 was restored to these cells by the expression of human *LUZP1-YFP* fusion (+LUZP1 cells). We examined LUZP1 localization associated with the actin cytoskeleton (Figure 5-figure supplement 1) and the centrosome (Figure 5-figure supplement 2) by immunofluorescence, and its levels by Western blot (Figure 5-figure supplement 3) in WT, Luzp1^−/−^ and +LUZP1 cells. WT, Luzp1^−/−^ and +LUZP1 cells were plated at equal densities and induced either to ciliate for 48 hours by serum withdrawal (starved), or to reabsorb their cilia by serum replenishment for 4 hours (refed) (Figure 5A). We quantified ciliation rates and primary cilia length at the mentioned timepoints. Luzp1^−/−^ fibroblasts displayed higher ciliation rate (60%) than WT (10.5%) and +LUZP1 (22.2%) when the cells were not subjected to starvation (Figure 5B). However, Luzp1^−/−^ cells were not significantly more ciliated than WT or +LUZP1 fibroblasts upon 48 hours of starvation or 4 hours after inducing cilia disassembly (Figure 5B). In addition, primary cilia in Luzp1^−/−^ cells were significantly longer than in non-starved WT cycling cells (Figure 5A,C); under starvation the differences among WT, Luzp1^−/−^ and +LUZP1 were not significant. Regarding cilia length, Luzp1^−/−^ and +LUZP1 cells behaved similarly (no starvation: WT 2.3 µm; Luzp1^−/−^ cells 3.0 µm; +LUZP1 cells 2.9 µm; 48 hours starvation: WT 4.2 µm; Luzp1^−/−^ cells 4.1 µm; +LUZP1 cells 4.8 µm; 4 hours after induction of disassembly: WT 2.4 µm; Luzp1^−/−^ cells 3.0 µm; +LUZP1 cells 2.9 µm; all average measures) (Figure 5A,C). These results confirm that Luzp1^−/−^ cells display longer and more abundant primary cilia compared to WT cells in cycling conditions and indicate that LUZP1 might affect primary cilia dynamics. One key event in ciliogenesis is the depletion of CCP110 and its partner CEP97 from the distal end of the MC, promoting the ciliary activating program in somatic cells (12, 38–41). Our previous work demonstrated that TBS cells displayed longer and more abundant cilia and that CCP110 underwent premature displacement from the MC in non-starved TBS cells (18). Since the ciliogenesis phenotype in Luzp1^−/−^ cells is reminiscent to the one described in TBS cells, we hypothesized that CCP110 might be also prematurely displaced from the centrosome in Luzp1^−/−^ cells. In order to test this hypothesis, we analyzed the centrosomal localization of CCP110 in WT and Luzp1^−/−^ cells by immunofluorescence. CCP110 was present at the centrosome in a higher proportion of WT cells (84%) than Luzp1^−/−^ cells (19%) (Figure 5D,E). This result suggests that the lack of LUZP1 might result in CCP110 displacement at the centrosome, leading to higher frequency of ciliogenesis in Luzp1^−/−^ cells.

**Figure 5.**
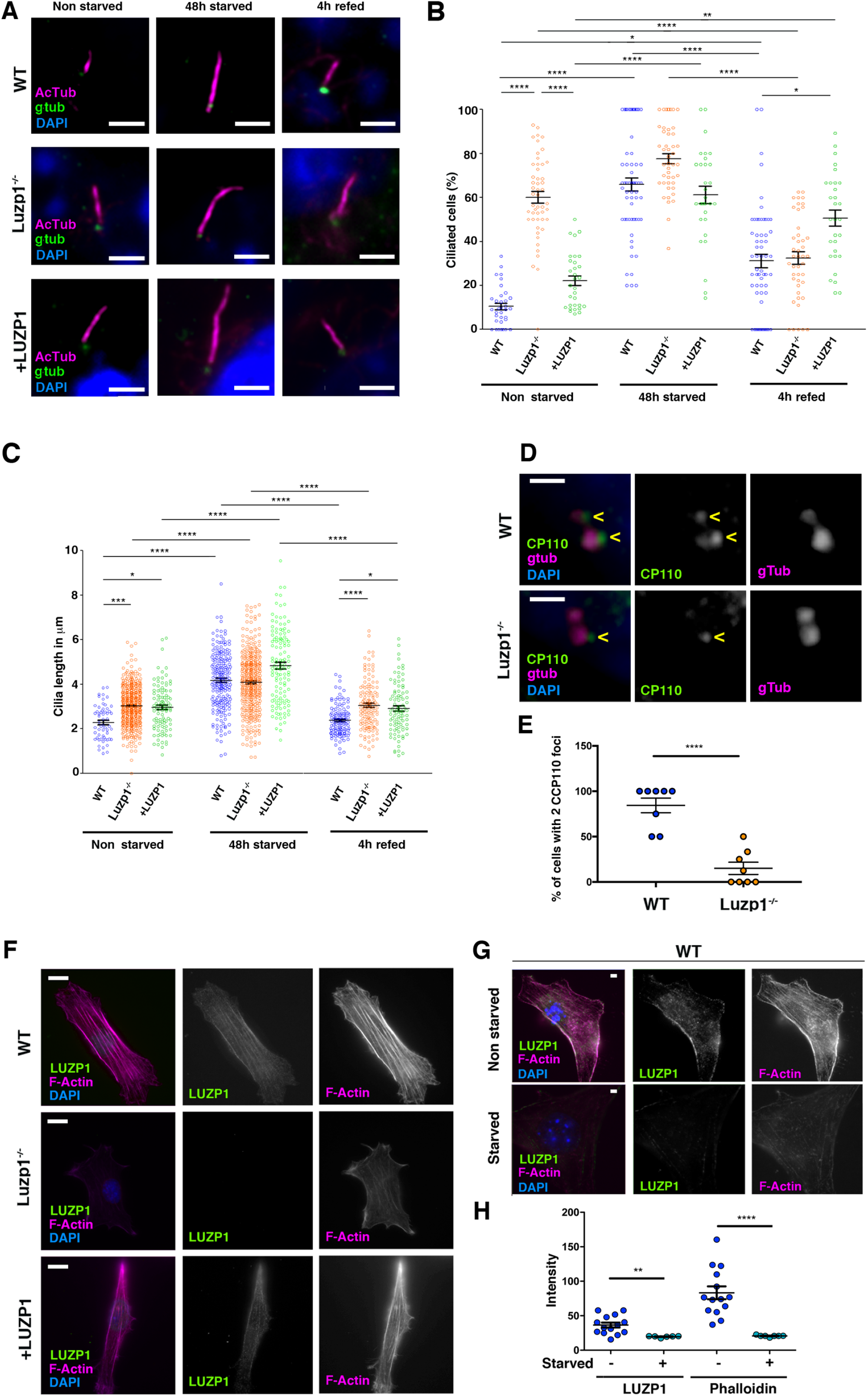
Luzp1^−/−^ cells show aberrant cilia frequency and length and reduced F-actin levels. (**A**) Micrographs of Shh-LIGHT2 cells (WT), Shh-LIGHT2 cells lacking *Luzp1* (Luzp1^−/−^) and Luzp1^−/−^ cells rescued with human *LUZP1-YFP* (*+*LUZP1) analyzed in cycling conditions (non-starved), or during cilia assembly (48 hours starved) and disassembly (4 hours re-fed). Cilia were visualized by acetylated alpha-tubulin (magenta), basal body by gamma-tubulin (green) and nuclei by DAPI (blue). Scale bar 2.5 µm. (**B, C**) Graphical representation of percentage of ciliated cells per micrograph (**B**) and cilia length (**C**) measured in WT (blue circles, n>34 micrographs), Luzp1^−/−^ (orange circles, n>44 micrographs) or *+*LUZP1 cells (green circles, n>30 micrographs) from three independent experiments. (**D**) Immunofluorescence micrographs of WT and LUZP1^−/−^ cells stained with antibodies against endogenous CCP110 (green), gamma tubulin to label the centrioles (purple) and DAPI to label the nuclei (blue). Black and white images show the single green and purple channels. Note the different distribution of CCP110 to the centrosome in LUZP1^−/−^ compared to WT cells. (**E**) Graphical representation of the percentage of cells showing the presence of CCP110 to both centrioles per micrograph corresponding to the experiments in (**D**); n=10 micrographs. Three independent experiments were pooled together. Pictures were taken using an Axioimager D1 fluorescence microscope, Zeiss, with a 63x objective. Scale bar, 1 µm. (**F**) Immunofluorescence micrographs of WT, Luzp1^−/−^ and *+*LUZP1 cells stained with an antibody against endogenous LUZP1 (green), phalloidin to detect F-actin (magenta), and counterstained with DAPI (blue). Single green and magenta channels are shown in black and white. Note the lack of LUZP1 in Luzp1^−/−^ cells. Scale bar, 10 µm. (**G**) Immunofluorescence micrographs of non-starved and starved WT cells stained with antibodies against endogenous LUZP1 (green), phalloidin (magenta) and DAPI (blue). Single green and magenta channels are shown in black and white. Scale bar, 5 µm. Imaging was performed using widefield fluorescence microscopy (Zeiss Axioimager D1, 63x objective). (**H**) Graphical representation of the LUZP1 or F-actin mean intensity as shown in (**G**). Graphs represent Mean and SEM of three independent experiments pooled together. P-values were calculated using One-way ANOVA and Bonferroni post-hoc test. The following figure supplement is available for Figure 5: Figure 5-figure supplement 1. LUZP1 mutant cells and antibody validation at the cytoskeleton. Figure 5-figure supplement 2. LUZP1 mutant cells and antibody validation at the centrosome. Figure 5-figure supplement 2. LUZP1 and antibody validation by Western blot.

Based on the LUZP1 localization to the actin cytoskeleton and that a reduction in LUZP1 was accompanied by a diminishment in F-actin levels in TBS^275^ cells, we hypothesised that LUZP1 might affect F-actin levels. First, we observed a reduction in F-actin (labelled by phalloidin) in the Luzp1^−/−^ cells compared to WT, which was recovered in +LUZP1 cells (Figure 5F). Furthermore, LUZP1 levels and actin filaments were diminished in WT fibroblasts upon starvation (Figure 5G,H). These results suggest that LUZP1 might stabilize actin and that starvation triggers both LUZP1 and F-actin reduction.

### Luzp1^−/−^ cells exhibit aberrant Sonic Hedgehog signaling

It is well-established that mammalian Shh signal transduction is dependent on functional primary cilia (42, 43). Therefore, we examined whether Shh signaling is compromised in Luzp1^−/−^ cells. Cells were starved for 24 hours and incubated in the presence or absence of purmorphamine (a SMO agonist) for 6 or 24 hours to activate the Shh pathway. The mRNA expression of two Shh target genes (*Gli1* and *Ptch1*) was quantified by qRT-PCR (Figure 6A,B). We found that the basal *Gli1* and *Ptch1* expression levels in Luzp1^−/−^ cells were higher than in WT cells (*Gli1* 1.5 fold and *Ptch1* 2.3 fold increase in Luzp1^−/−^ *vs* WT cells without purmorphamine) (Figure 6A,6B). Upon induction by purmorphamine for 24 hours, WT cells increased significantly the expression of both targets, while Luzp1^−/−^ cells did not, indicating that Luzp1^−/−^ cells fail to induce Shh signaling. To further study the role of LUZP1 in Shh signaling, we analyzed GLI3 processing by Western blot using total lysates extracted from WT *vs* Luzp1^−/−^ cells. Without purmorphamine induction, we found a significantly higher ratio of GLI3 activating form *vs* GLI3 repressive form (GLI3-A:GLI3-R) in Luzp1^−/−^ cells compared to WT (2.9 fold increase in Luzp1^−/−^ cells *vs* WT) (Figure 6C and Figure 6-figure supplement 1). After induction, the values were similar for Luzp1^−/−^ and WT cells. Since the Luzp1^−/−^ parental line was Shh-LIGHT2, we also examined the effects of lacking *Luzp1* on Shh signaling by measuring the activity of a GLI-responsive Firefly luciferase reporter (Figure 6D). Prior to purmorphamine treatment, Luzp1^−/−^ cells showed higher Shh activity compared to control or +LUZP1 cells, as observed in TBS-derived cells (1.6 fold-activity in Luzp1^−/−^ *vs* 0.5 fold-activity in +LUZP1 cells or 1 fold-activity in WT cells) (18). However, the induction capacity of Luzp1^−/−^ cells upon purmorphamine treatment was reduced compared to WT. Altogether, the observed defects in *Ptch1* and *Gli1* gene expression, reduced GLI3 processing and Shh reporter misregulation confirm a role for LUZP1 in Shh signaling.

**Figure 6.**
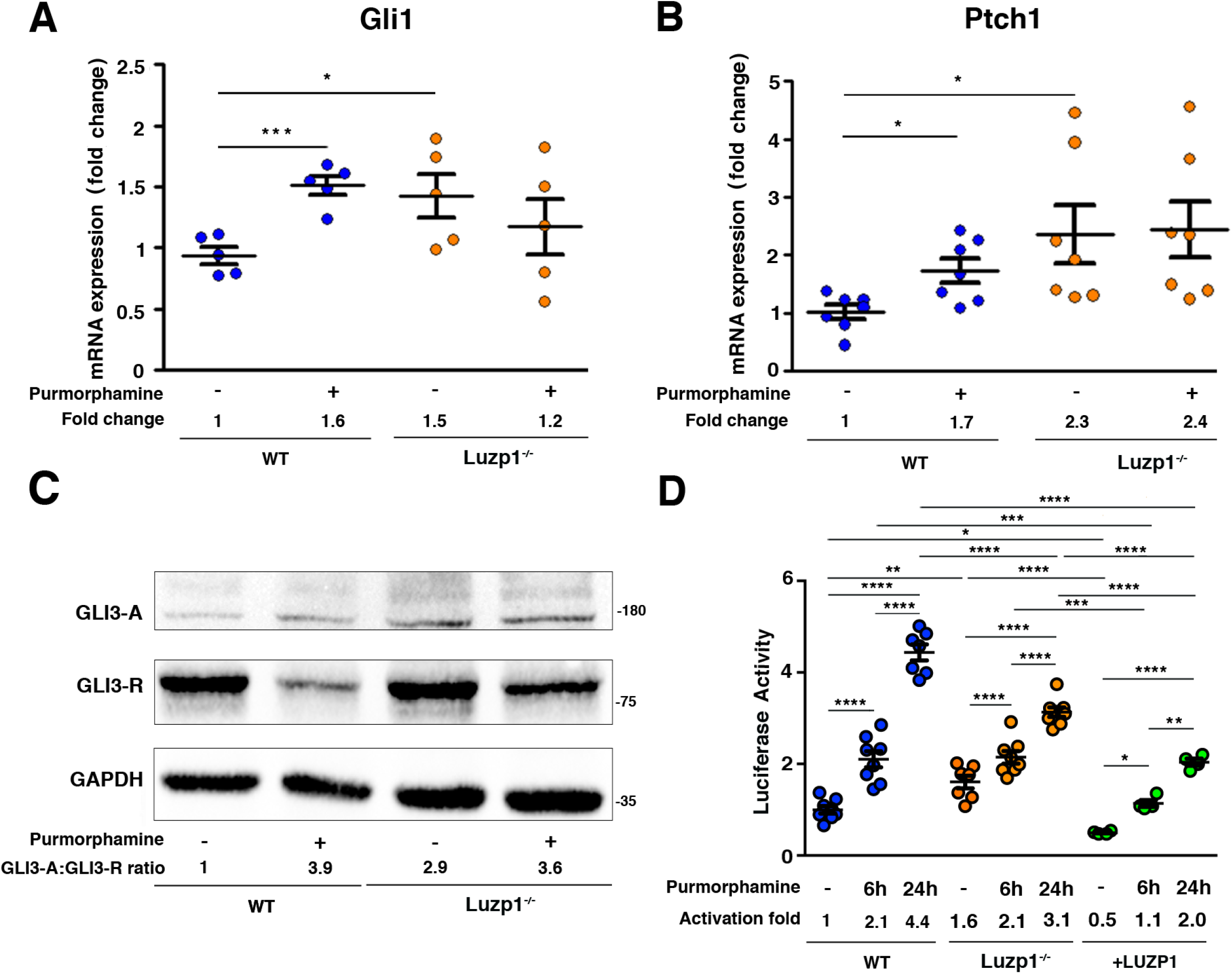
Luzp1^−/−^ cells show aberrant Shh signaling. (**A**, **B**) Graphical representation of the fold-change in the expression of *Gli1* (n=5) (**A**) and *Ptch1* (n=7) (**B**) obtained by qPCR from wild-type Shh-LIGHT2 cells (WT; blue dots) or Shh-LIGHT2 cells lacking *Luzp1* (Luzp1^−/−^; orange dots), treated (+) or not (-) with purmorphamine for 24 hours. **(C)** Western blot analysis of lysates from WT and Luzp1^−/−^ cells. Samples were probed against GLI3 that detects both GLI3-activator form (GLI3-A) and GLI3-repressor form (GLI3-R) and GAPDH was used as loading control. Numbers under the lanes are the result of dividing the activator by the repressor intensities, taking WT non-induced value as 1. Molecular weight markers are shown to the right. (**D**) Graphical representation of fold-change in luciferase activation when WT (n>7; blue dots), Luzp1^−/−^ (n>7; orange dots) or *+*LUZP1 (n=4; green dots) cells are treated for 6 and 24 hours or not (-) with purmorphamine. All graphs represent the Mean and SEM. P-values were calculated using two-tailed unpaired Student’s t-test or One-way ANOVA and Bonferroni post-hoc test. The following figure supplement is available for Figure 6: Figure 6-figure supplement 1. Western blot full pictures for Figure 6.

### Truncated SALL1 promotes LUZP1 degradation through the ubiquitin proteasome system (UPS) pathway

In concordance with immunofluorescence results in Figures 3B and 4A, we confirmed a reduction in total LUZP1 levels in TBS^275^ cells compared to controls by Western blot (Figure 7A,B and Figure 7-figure supplement 1). Because no transcriptional changes in *LUZP1* expression were detected between control and TBS^275^ samples (Figure 7-figure supplement 2), we hypothesized that truncated SALL1 might lead to ubiquitin-proteasome system (UPS)-mediated LUZP1 degradation. We therefore analyzed LUZP1 levels after treatment with the proteasomal inhibitor MG132, both in control and TBS^275^ cells. LUZP1 levels were increased to a higher extent in TBS^275^ compared to control cells (1.8 fold increase in control *vs* 2.4 fold increase in TBS^275^ cells) (Figure 7A,B). Moreover, we confirmed the reduction of LUZP1 levels in the CRISPR/Cas9 TBS model cell line (293^335^), in which the SALL1 hot-spot region was mutated, compared to its parental cell line (293^WT^) (Figure 7C,D and Figure 7-figure supplement 1), and likewise in HEK 293FT cells overexpressing truncated SALL1 (*SALL1^275^-YFP*) compared to cells overexpressing *YFP* as control (Figure 7E,F and Figure 7-figure supplement 1). A more prominent increase in LUZP1 accumulation upon MG132 treatment was also observed in 293^335^ and HEK 293FT cells overexpressing *SALL1^275^-YFP* compared to controls (Figure 7C,D and Figure 7E,F, respectively, and Figure7-figure supplement 1). Additionally, we also observed LUZP1 accumulation upon MG132 treatment by immunofluorescence in RPE1 cells, both at the actin cytoskeleton (Figure 7G, upper panels) and at the centrosome (Figure 7G, lower panels). All together, these results show that LUZP1 levels are sensitive to degradation via the UPS pathway and suggest that truncated SALL1 may contribute to this process. Furthermore, we compared LUZP1 ubiquitination in 293^WT^ *vs* 293^335^ cells using the BioUb strategy (see Materials and Methods) (44). We could observe a prominent band in presence of BioUb, likely corresponding to the monoubiquitinated form of LUZP1 in the pulldowns (Figure 7G). This form was present in 293^WT^ and 293^335^ cells, and increased in both cases in presence of MG132. In addition, we observed a smear at higher molecular weight corresponding to polyubiquitinated forms of LUZP1 (Figure 7G, Biotin PD). Notably, the LUZP1 ubiquitinated pool relative to the input levels was higher in 293^335^ compared to 293^WT^ cells upon MG132 treatment (Figure 7G, Biotin PD, lane 8 *vs* lane 11). These results reflect that truncated SALL1 promotes LUZP1 degradation through the UPS pathway.

**Figure 7.**
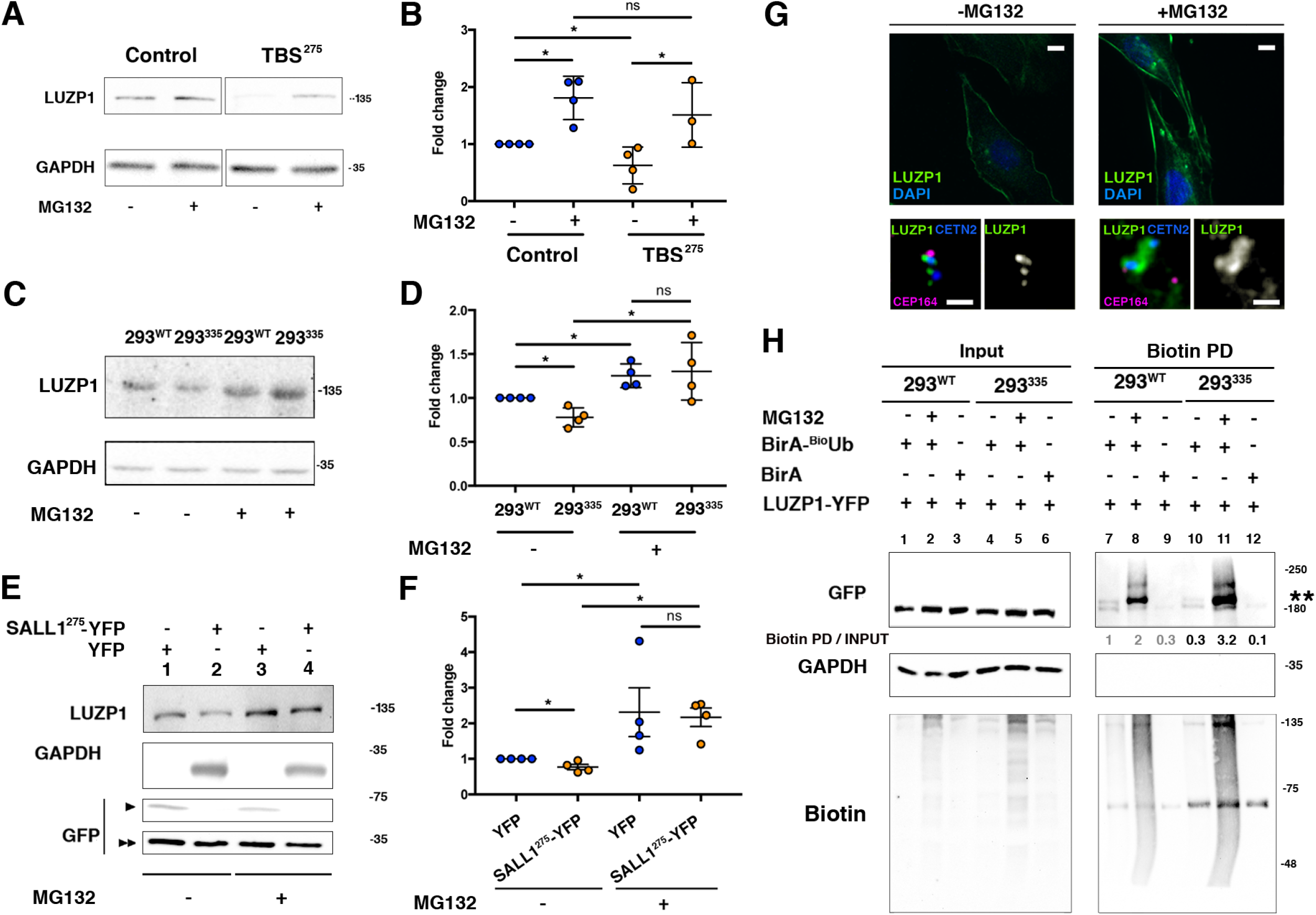
Truncated SALL1 leads to LUZP1 degradation through the UPS. (**A**) Representative Western blot of ESCTRL2 and TBS^275^ total cell lysates treated or not with MG132. A specific antibody detected endogenous LUZP1, and GAPDH was used as loading control. (**B**) Graphical representation of the fold changes of LUZP1/GAPDH ratios obtained in (**A**) for of ESCTRL2 (blue dots) and TBS^275^ (orange dots) treated (+) or not (-) with the proteasome inhibitor MG132. Note the increase of LUZP1 until reaching control levels in TBS^275^ cells upon MG132 treatment. (**C**) Representative Western blot of 293^WT^ and 293^335^ total cell lysates treated or not with MG132. A specific antibody against LUZP1 detected endogenous LUZP1, and GAPDH was used as loading control. (**D**) Graphical representation of the fold changes of LUZP1/GAPDH ratios obtained in panel C for 293^WT^ (blue dots) and 293^335^ (orange dots) treated (+) or not (-) with MG132. Note that LUZP1 in 293^335^ reaches control levels with MG132 treatment. (**E**) Representative Western blot of total lysates of HEK 293FT cells transfected with *SALL1^275^-YFP* (lanes 1 and 3) or *YFP* alone (lanes 2 and 4) treated (+) or not (-) with MG132. Specific antibodies against LUZP1, GFP and GAPDH were used. (**F**) Graphical representation of the fold changes of LUZP1/GAPDH ratios obtained in (**E**) for HEK 293FT cells transfected with *SALL1^275^-YFP* (orange dots) or *YFP* alone (blue dots) treated (+) or not (-) with MG132. Note that LUZP1 increases in the presence of MG132 when *SALL1^275^-YFP* was transfected. Data from at least three independent experiments pooled together are shown. P-values were calculated using two-tailed unpaired Student’s t-test. (**G**) Immunofluorescence micrographs of RPE1 cells treated (+MG132) or not (-MG132) with proteasome inhibitor showing LUZP1 in the cytoskeleton (upper panels) or in the centrosome (lower panels). Note the overall increase of LUZP1 upon MG132 treatment. Scale bar 10 µm (cytoskeleton panels) or 0.5 µm (centrosome panels). Images were taken using widefield fluorescence microscopy (Zeiss Axioimager D1, 63x objective). (**H**) Western blot analysis of input and biotin pulldown (PD) of HEK 293FT cells transfected with *LUZP1-YFP* and BioUb or BirA alone treated (+) or not (-) with MG132. Specific antibodies (GFP, GAPDH, Biotin) were used as indicated. Numbers under GFP panel are the result of dividing each biotin PD band intensity by the respective input band intensity and normalize them to lane 1. Molecular weight markers in kDa are shown to the right. Two asterisks indicate monoubiquitinated LUZP1. The following figure supplement is available for Figure 7: Figure 7-figure supplement 1. Western blot full pictures for Figure 7. Figure 7-figure supplement 2. LUZP1 mRNA expression levels.

### LUZP1 overexpression represses cilia formation and increases F-actin levels in human fibroblasts

Our results suggest that LUZP1 could be a mediator of TBS cilia phenotype and that this could be caused, at least in part by the increased degradation of LUZP1 triggered by truncated SALL1. Therefore, increasing LUZP1 levels in TBS cells might affect the cilia and actin cytoskeleton phenotypes. To check whether *LUZP1* overexpression is sufficient to repress ciliogenesis in primary human fibroblasts, Control and TBS^275^ cells were transduced with *YFP* or *LUZP1-YFP* (Figure 8A). Whereas most non-transduced surrounding cells, as well as 100 % of the TBS^275^ cells expressing *YFP* were ciliated, only 40% of the Control and TBS^275^ cells transduced with *LUZP1-YFP* displayed cilia (Figure 8B). Furthermore, we aimed to rescue the actin cytoskeleton defects observed in TBS^275^ cells by overexpressing *LUZP1-YFP*. Immunostaining showed that LUZP1-YFP overexpression led to an increase in F-actin levels both in control and in TBS^275^ cells compared to the surrounding non-transfected cells or TBS^275^ cells overexpressing *YFP* (Figure 8C). All together, these results support the notion that LUZP1 may be a potential negative regulator of cilia formation and an F-actin stabilizing protein.

**Figure 8.**
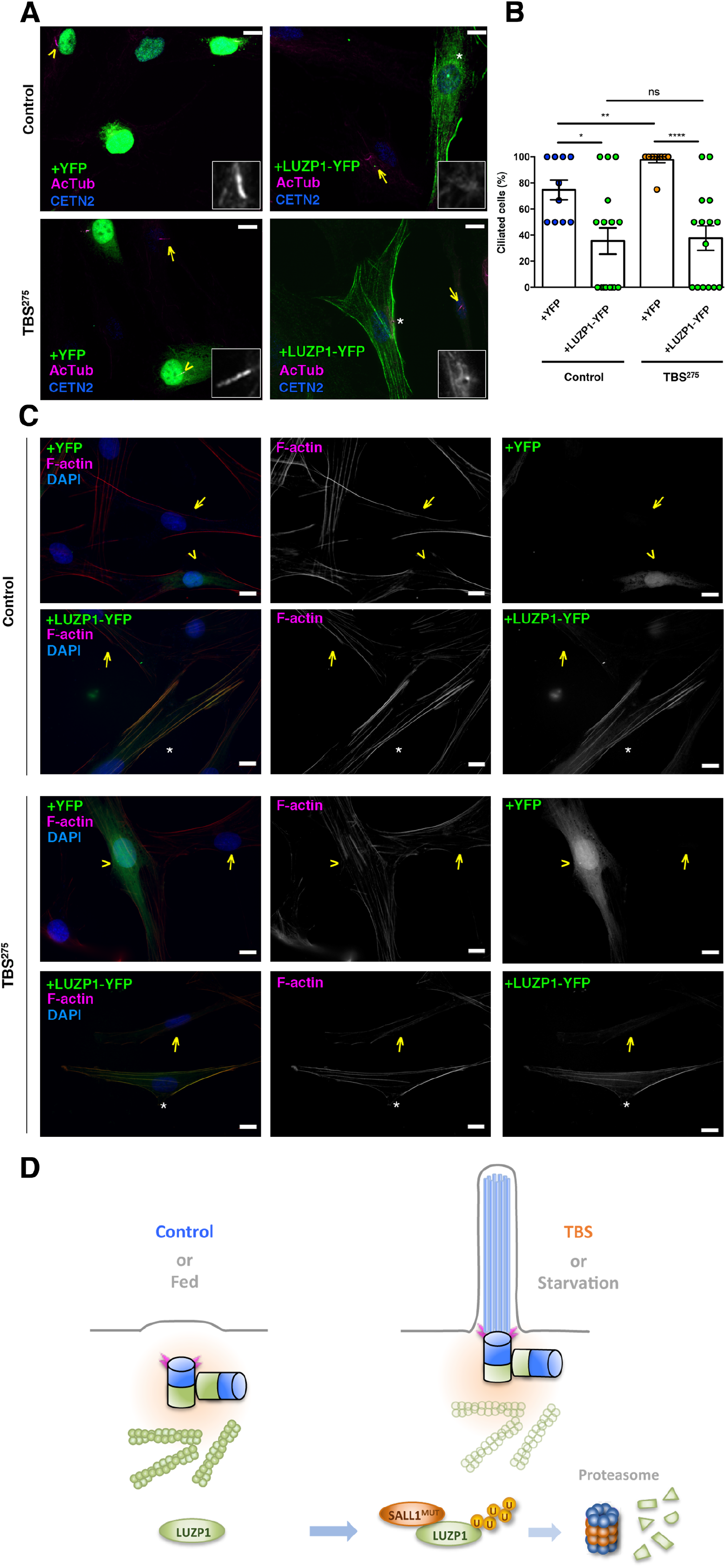
LUZP1 overexpression suppresses ciliogenesis and increases F-actin levels. (**A**) Representative micrographs of ciliated control and TBS^275^ cells overexpressing *YFP* or *LUZP1-YFP*. Yellow arrowhead and white asterisk point at a magnified region shown in the lower right panel in black and white. Note the lack of cilia in cells overexpressing LUZP1-YFP (white asterisk) compared to non-transfected cells (yellow arrow). AcTub: acetylated alpha tubulin; CETN2: Centrin-2. (**B**) Graphical representation of the number of ciliated cells per micrograph in control and TBS^275^ cells overexpressing *YFP* or *LUZP1-YFP* (n>10 micrographs). Graphs represent Mean and SEM of ciliation frequencies per micrograph in three independent experiments pulled together. P-values were calculated using One-way ANOVA and Bonferroni post-hoc test or two-tailed unpaired Student’s t-test. (**C**) Representative micrographs of Control and TBS^275^ cells overexpressing *YFP* (yellow arrowhead) or *LUZP1-YFP* (white asterisk) co-stained with phalloidin to label F-actin and DAPI. Note the increase in F-actin levels in cells overexpressing LUZP1-YFP (white asterisk) compared to non-transfected cells (yellow arrow). Scale bar, 10 µm. Imaging was performed using widefield fluorescence microscopy (Zeiss Axioimager D1, 63x objective). (**D**) A model representing how the presence of truncated SALL1 underlies cilia and actin malformations in TBS through LUZP1 interaction and UPS-mediated degradation. In control (or fed) cells (left), LUZP1 (in green) localizes to F-actin and to the proximal ends of the two centrioles, inhibiting cilia formation. By contrast, in TBS (or starved) cells (right) the truncated form of SALL1 interacts with LUZP1, leading to its UPS-mediated degradation. As a result, LUZP1 levels are reduced both at the centrosome and at the cytoskeleton, which will allow the formation of the primary cilia.

## DISCUSSION

Our results indicate that LUZP1 might be a mediator of the TBS phenotype via its interaction with truncated SALL1 and its effect on mammalian ciliogenesis: i) LUZP1 localization is altered both at the centrosome and actin cytoskeleton in TBS-derived cells; ii) LUZP1 levels are reduced in TBS-derived cells likely due to truncated SALL1-mediated degradation through the UPS; iii) LUZP1 interacts with important regulators of ciliogenesis (CCP110, CEP97) and of the actin cytoskeleton (FLNA); and iv) in the absence of LUZP1, the assembly and growth of primary cilia is enhanced in cycling cells, accompanied by an increase in basal Shh signaling. Our findings uncover a perturbation of cilia and actin cytoskeleton in the absence of LUZP1. Cells adapt to serum starvation, i.e. a reduction in nutrients and growth factors, by coordinated cytoskeletal rearrangements and cilia signaling. This integrated response requires signal transduction relays that communicate the cytoplasmic actin polymerization status with cilia. Here, we propose that LUZP1 might act as a nexus in this complex intracellular network and that truncated SALL1 disrupts this network.

### LUZP1 localizes to the centrosome and actin cytoskeleton

LUZP1 was previously described as a nuclear protein, with expression limited to the mouse brain (28, 29). We tested two different commercial antibodies against LUZP1 and, while nuclear localization was weakly detected by immunofluorescence, we observed a more prominent localization of LUZP1 to the actin cytoskeleton and centrosome, both in human and mouse cells. This localization is consistent with our TurboID analysis that showed an enrichment of factors associated with the actin cytoskeleton and/or centrosomes among the potential interactors of LUZP1. The localization of LUZP1 to the actin cytoskeleton, as well as being expressed in tissues beyond the brain, is consistent with independent validation in cell lines by the Human Protein Atlas (HPA; proteinatlas.org) and other expression databases (e.g. EMBL EBI Expression Atlas ebi.ac.uk/gxa). Moreover, two independent proximity labeling studies identified LUZP1 as a proximal interactor of centriole (35) and centriolar satellite-related proteins (45). Here, we report that LUZP1 forms a basket-like 3D structure surrounding the proximal end of both centrioles. Like LUZP1, a large number of centrosomal scaffold proteins (as for instance Cep120, Cep57, Cep63, Cep152, CPAP, Cdk5Rap2, PCNT, among others) contain coiled-coil regions, and the proteins are concentrically localized around a centriole in a highly organized fashion (5–7). Furthermore, here we show that LUZP1 interacts with centrosome and actin-related proteins (Figure 3 and Figure 4). LUZP1 has also been identified as an interactor of ACTR2 (ARP2 actin related protein 2 homologue) and FLNA (19, 20), and it has been recently described as an actin cross-linking protein (19). Additionally, we found that LUZP1 localizes not only to centrioles and actin cytoskeleton, but also to the midbody in dividing cells, which was recently reported to influence ciliogenesis in polarised epithelial cells (46).

Discrepancies with the previously reported LUZP1 localization and distribution might be due to technical differences, or perhaps the epitope specificity for the previously reported antiserum. Our data suggest that the association of LUZP1 to centrosomes and actin filaments in many tissues may contribute to its overall roles.

### LUZP1 is altered in TBS-derived cells

TBS is caused by mutations in *SALL1* gene, which lead to the formation of a truncated protein that interferes with the normal function of the cell. Here we found that LUZP1interacts with truncated SALL1 and with SALL1^FL^, which suggests that interaction occurs through an N-terminal domain shared by both. We believe that, in control cells, LUZP1 and SALL1^FL^ might have minimal interaction due to their respective localizations to the cytoplasm and nucleus. However, truncated SALL1, alone or together with SALL1^FL^ that is retained in the cytoplasm, likely interacts with cytoplasmic LUZP1, promoting its degradation and functional inhibition. Importantly, we detected an increase in LUZP1 levels upon treatment with the proteasome inhibitor MG132 (Figure 7), suggesting that LUZP1 degradation is proteasome-mediated. Next, we demonstrated that LUZP1 is ubiquitinated, and that truncated SALL1 both increases LUZP1 ubiquitination and decreases its stability. LUZP1 ubiquitination was detected in several proteomic screens for ubiquitinated proteins (47–51). The mechanism by which truncated SALL1 can influence LUZP1 ubiquitination is yet to be revealed, but one possibility could be *de novo* complexes involving specific Ub E3 ligases or de-ubiquitinases which could influence LUZP1 stability. In fact, various E3s/de-ubiquitinases were found as proximal interactors of truncated SALL1 and LUZP1, as well as other components of the UPS. Furthermore, regulation by the UPS system has been reported for centrosomal factors, necessary for the process of ciliogenesis such as CCP110 (52, 53, 54).

The phenotypes observed in TBS individuals fall within the spectrum of those observed in ciliopathies, characterized by malformations in digits, ears, heart, brain and kidneys. Defective regulation of cilia function and/or formation is a contributing factor in TBS (18). Both Luzp1^−/−^ and TBS cells showed a reduction in F-actin accompanied by an increase in ciliation. We suggest that the reduction in filamentous actin in TBS cells might contribute to their higher cilia abundance, longer cilia and increased Shh signaling. By increasing LUZP1 expression in control and TBS^275^ cells, F-actin levels are increased and cilia frequency is reduced, suggesting that LUZP1 may have a role in the TBS phenotype.

### LUZP1 as an integrator of actin and primary-cilium dynamics

Actin dynamics coordinate several processes that are crucial for ciliogenesis. For example, placing the MC to the appropriate area at the cell cortex is an actin-dependent process (55, 56). A reduction in cortical actin might potentially promote ciliogenesis, as there would be no physical restriction to prevent cilium growth. Supporting this hypothesis, several studies have found that changes in the actin network architecture, induced either chemically or genetically, promote ciliogenesis or affect cilia length (57–62). How actin regulates cilium length is not clear. One hypothesis is that actin is involved in ectocytosis and cilium tip scission, preventing the axoneme from growing too long (63, 64). Moreover, the removal of the CCP110/CEP97 complex from the centrosome is thought to be an essential event at the beginning of cilia formation. Many proteins are known to interact with the CCP110/CEP97 complex to regulate ciliogenesis (65). We found that LUZP1 is associated with CCP110 and CEP97 and that CCP110 was displaced in Luzp1^−/−^ cells. However, our TurboID analysis did not detect CCP110 and CEP97 in the vicinity of LUZP1. This divergence might result from the limitation of TurboID to detect proteins that are separated further than 10 nm from each other. In fact, we found LUZP1 and CCP110 localizing to de proximal and distal end of centrioles, respectively (Figure 2D).

### The role of LUZP1 in neural tube closure and cardiac defects

Several studies have emphasized the tight links between cytoskeletal organization and cell fate and have implicated Shh signaling in the etiology of neural tube closure defects (66). Shh signaling is aberrant in TBS patient-derived fibroblasts (18). While reporters of Shh signaling were not examined in Luzp1 KO mice, there was ectopic Shh expression reported in the neuroepithelium of the *Luzp1* KO mouse hindbrain, which displays NTD (23). Here, we show aberrant Shh signaling in Luzp1^−/−^ cells. Our results might indicate that LUZP1 is therefore pivotal to Shh signaling such that cranial neural tube closure may be achieved. In addition, Hsu *et. al.* noted that, in the *Luzp1 KO* embryos, exencephaly may be caused by failure in bending at the dorsolateral hinge point and that the dorsolateral neural folds were convex instead of the concave morphology observed in WT embryos (23). It has been reported that changes in apical actin architecture are required for the proper formation of the neural tube (67). Thus, we hypothesize that actin defects may contribute to neural tube defects observed in *Luzp1* KO mice. Likewise, aberrant primary cilia and Shh signaling might be present in those mice, both of which are known to interfere with neural tube closure (68).

In addition to NTDs, *Luzp1* knockout mice phenocopy another feature often associated with human ciliopathies, namely cardiac malformation, which can also occur in TBS patients (69). TBS cardiac defects include atrial or ventricular septal defect, the latter of which is seen in *Luzp1* knockout mice. Moreover, compound *Sall1/Sall4* KO mutant mice exhibit both NTDs and cardiac problems (70). While Luzp1 and Sall1 may both contribute to brain and heart development, a novel crosstalk may arise in TBS due to dominantly-acting truncated SALL1 that can derail these processes and cause deformities.

In conclusion, our data indicate that LUZP1 localizes to actin stress fibers and to the centrosome, where it may act as a cilia suppressor (Figure 8D). Upon starvation, overall LUZP1 levels are diminished in both structures, which facilitates the formation of the primary cilia. Starved control cells appear similar to fed TBS cells, in which a truncated form of SALL1 localizes to the cytoplasm, interacting with LUZP1 and enhancing its degradation. As a result, the frequency of cilia formation increases, and cilia are longer than in control cells. Our findings point to the intriguing possibility that LUZP1 might be a key relay switch between the actin cytoskeleton and cilia regulation and along with other factors, might contribute to the phenotypes observed in TBS.

## MATERIALS AND METHODS

### Cell culture

TBS-derived primary fibroblasts, U2OS, HEK 293FT (Invitrogen), and mouse Shh-LIGHT2 cells (37) were cultured at 37°C and 5% CO_2_ in Dulbecco’s modified Eagle medium (DMEM) supplemented with 10% foetal bovine serum (FBS, Gibco) and 1% penicillin/streptomycin (Gibco). Human telomerase reverse transcriptase immortalized retinal pigment epithelial cells (TERT-RPE1, ATCC CRL-4000) were cultured in DMEM:F12 (Gibco) supplemented with 10% FBS and 1% penicillin and streptomycin. Dermal fibroblasts carrying the *SALL1* pathogenic variant c.826C>T (*SALL1^c.826C>T^*), that produce a truncated protein p.Leu275* (SALL1^275^), were derived from a male TBS individual UKTBS#3 (called here TBS^275^) (18). Adult female dermal fibroblasts (ESCTRL#2) from healthy donors were used as controls. Cultured cells were maintained between 10 and 20 passages, tested for senescence by γ-H2AX staining, and grown until confluence (6-well plates for RNA extraction and Western blot assays; 10 cm dishes for pulldowns). The use of human samples in this study was approved by the institutional review board (Ethics Committee at CIC bioGUNE) and appropriate informed consent was obtained from human subjects or their parents.

### Cell synchronization and drug treatment

hTERT RPE-1 cells (RPE-1) were arrested in G1 phase by treatment with mimosine (Sigma, 400 µM) for 24 hours. For S phase arrest, cells were subjected to thymidine treatment (Sigma, 2.5 mM) for 16 hours, followed by release for 8 hours, and subsequently blocked again for 16 hours. For G2/M phase arrest, cells were treated with RO-3306 (Sigma, 10 µM) for 20 hours. For entering G0 phase and induce primary cilia formation, cells were starved for 24 hours (DMEM, 0% FBS, 1% penicillin and streptomycin). Cells were treated with the proteasome inhibitor MG132 (Calbiochem, 5 µM) for 15 hours and with Cytochalasin D (Sigma, 10 µM) for 10 minutes to stimulate actin depolymerization. HEK 293FT cells were transfected using calcium phosphate method and U2OS cells using Effectene Transfection Reagent (Qiagen). To induce primary cilia, cells were starved for at least 24 hours (DMEM, 0% FBS, 1% penicillin and streptomycin).

### CRISPR-Cas9 genome editing

CRISPR-Cas9 targeting of *SALL1* locus was performed to generate a HEK 293FT cell line carrying a TBS-like allele (18). The mouse *Luzp1* locus was targeted in NIH3T3-based Shh-LIGHT2 fibroblasts (37) (kind gift of A. McGee, Imperial College). These are NIH3T3 mouse fibroblasts that carry an incorporated Shh reporter (firefly luciferase under control of *Gli3*-responsive promoter). Cas9 was introduced into Shh-LIGHT2 cells by lentiviral transduction (Lenti-Cas9-blast; Addgene #52962; kind gift of F. Zhang, MIT) and selection with blasticidin (5 µg/ml). Two high-scoring sgRNAs were selected (http://crispr.mit.edu/) to target near the initiation codon (sg2: 5’-CTTAAATCGCAGGTGGCGGT_TGG-3’; sg3: 5’-CTTCAATCTTCAGTACCCGC_TGG-3’). These sequences were cloned into px459 2.0 (Addgene #62988; kind gift of F. Zhang, MIT), for expressing both sgRNAs and additional Cas9 with puromycin selection. Transfections were performed in Shh-LIGHT2/Cas9 cells with Lipofectamine 3000 (Thermo). 24 hours after transfection, transient puromycin selection (0.5 µg/ml) was applied for 48 hours to enrich for transfected cells. Cells were plated at clonal density, and well-isolated clones were picked and propagated individually. Western blotting was used to identify clones lacking *Luzp1* expression. Further propagation of a selected clone (#6) was carried out with G418 (0.4 mg/ml) and zeocin (0.15 mg/ml) selection to maintain expression of luciferase reporters. Genotyping was performed using genomic PCR (*MmLuzp1_geno_for*: 5’-GTTGCCAAAGAAGGTTGTGGATGCC-3’; *MmLuzp1_geno_rev*: 5’-CGTAAGGTTTTCTTCCTCTTCAAGTTTCTC-3’) and revealed a homozygous deletion of bases between the two sgRNA target sites, predicting a frame-shifted truncated protein (MAELTNYKDAASNRY*), and resulting in a null *Luzp1* allele. A rescue cell line was generated by transducing Shh-LIGHT2 *Luzp1* KO clone #6 with a lentiviral expression vector carrying EFS-LUZP1-YFP-P2A-blast^R^, with a positive population selected by fluorescence-activated cell sorting.

### Plasmid construction

*SALL1* truncated (*SALL1^275^-YFP or Myc-BirA*-SALL1^275^*) and FL versions (*SALL1^FL^-YFP*, *SALL1^FL^-2xHA or Myc-BirA*-SALL1^FL^)* were previously described (18). To identify bio-ubiquitin conjugates, human *LUZP1* ORF was amplified by high-fidelity PCR (Platinum SuperFi; Thermo) from hTERT-RPE1 cDNA and cloned to generate *CB6-GFP-LUZP1*. This was used as a source clone to generate additional variants (*CMV-LUZP1-YFP*, *Myc-TurboID-LUZP1*). The LUZP1-YFP and TurboID-LUZP1 lentiviral expression vectors were generated by replacing Cas9 in Lenti-Cas9-blast (Addgene #52962). All constructs were verified by Sanger sequencing. Plasmids *CAG-BioUBC(x4)_BirA_V5_puro* (called here BioUb) and *CAG-BirA-puro* (called here BirA) were reported previously (44).

### Biotin pulldowns

Using the BioID and the TurboID methods (33, 71), proteins in close proximity to SALL1 and LUZP1, respectively, were biotinylated and isolated by streptavidin-bead pulldowns. *Myc-BirA*-SALL1^c.826C>T^*, *Myc-BirA*-SALL1^FL^* or *Myc–TurboID-LUZP1* were transfected in HEK 293FT cells (10 cm dishes). For the isolation of BioUb-conjugates 10 cm dishes were transfected with BioUb or BirA as control, according to (Pirone et al 2016). Briefly, 24 hours after transfection, medium was supplemented with biotin at 50 µM. Cells were collected 48 hours after transfection, washed 3 times on ice with cold phosphate buffered saline (PBS) and scraped in lysis buffer [8 M urea, 1% SDS, 1x protease inhibitor cocktail (Roche), 60 µM NEM in 1x PBS; 1 ml per 10 cm dish]. At room temperature, samples were sonicated and cleared by centrifugation. Cell lysates were incubated overnight with 40 μl of equilibrated NeutrAvidin-agarose beads (Thermo Scientific). Beads were subjected to stringent washes using the following washing buffers (WB), all prepared in PBS: WB1 (8 M urea, 0.25% SDS); WB2 (6 M Guanidine-HCl); WB3 (6.4 M urea, 1 M NaCl, 0.2% SDS), WB4 (4 M urea, 1 M NaCl, 10% isopropanol, 10% ethanol and 0.2% SDS); WB5 (8 M urea, 1% SDS); and WB6 (2% SDS). For elution of biotinylated proteins, beads were heated at 99°C in 50 μl of Elution Buffer (4x Laemmli buffer, 100 mM DTT). Beads were separated by centrifugation (18000 x g, 5 minutes).

### Lentiviral transduction

Lentiviral expression constructs were packaged using psPAX2 and pVSV-G (Addgene) in HEK 293FT cells, and lentiviral supernatants were used to transduce Shh-LIGHT2 cells. Stable-expressing populations were selected using puromycin (1 µg/ml) or blasticidin (5 µg/ml). Lentiviral supernatants were concentrated 100-fold before use (Lenti-X concentrator, Clontech). Concentrated virus was used for transducing primary fibroblasts and hTERT-RPE1 cells.

### Mass spectrometry

Analysis was done in hTERT-RPE1 cells stably expressing TurboID-LUZP1 at near endogenous levels. Three independent pulldown experiments (1×10^7^ cells per replicate) were analyzed by MS. Samples eluted from the NeutrAvidin beads were separated in SDS-PAGE (50% loaded) and stained with Sypro-Ruby (Biorad) according to manufacturer’s instructions. Entire gel lanes were excised, divided into pieces and in-gel digested with trypsin. Recovered peptides were desalted using stage-tip C18 microcolumns (Zip-tip, Millipore) and resuspended in 0.1% FA prior to MS analysis. In this study, samples (33%) were loaded onto a nanoElute liquid chromatograph (Bruker) at 300 nl/min and using a 15 min linear gradient of 3–45% acetonitrile, coupled on-line to a TIMS TOF Pro mass spectrometer using PASEF (Bruker Daltonics) (72). Data was processed by Data Analysis v3.0 (Bruker) and searches were carried out by Mascot (MatrixScience). Applied search parameters were: 50 ppm precursor ion tolerance and 0.05 Da for fragment ions; Carbamidomethylation as fixed and methionine oxidation as variable modifications; up to two missed cleavages. Database search was performed against UNIPROT database (December 2018) containing only *Homo sapiens* entries.

Protein IDs were ranked according to the number of peptides found and their corresponding intensities. Gene ontology (GO) term enrichment was analyzed using g:GOSt Profiler, a tool integrated in the g:Profiler web server (73). GO enrichment was obtained by calculating –log_10_ of the P-value.

### GFP-trap pulldowns

All steps were performed at 4°C. HEK 293FT transfected cells were collected after 48 hours, washed 3 times with 1x PBS and lysed in 1 ml of lysis buffer [25 mM Tris-HCl pH 7.5, 150 mM NaCl, 1 mM EDTA, 1% NP-40, 0.5% Triton X-100, 5% glycerol, protease inhibitors (Roche)]. Lysates were kept on ice for 30 minutes vortexing every 5 minutes and spun down at 25,000 x g for 20 minutes. After saving 40 µl of supernatant (input), the rest was incubated overnight with 30 µl of pre-washed GFP-Trap resin (Chromotek) in a rotating wheel. Beads were washed 5 times for 5 minutes each with WB (25 mM Tris-HCl pH 7.5, 300 mM NaCl, 1 mM EDTA, 1% NP-40, 0.5% Triton X-100, 5% glycerol). Beads were centrifuged at 2,000 x g for 2 minutes after each wash. For elution, samples were boiled for 5 minutes at 95°C in 2x Laemmli buffer.

### Immunoprecipitation

All steps were performed at 4°C. Cells were collected and lysates were processed as described for GFP-trap pulldowns. After saving 40 µl of supernatant (input), the rest was incubated overnight with 1 µg of anti-CEP97 antibody (Proteintech), or anti-LUZP1 antibody (Sigma) and for additional 4 hours with 40 µl of pre-washed Protein G Sepharose 4 Fast Flow beads (GE Healthcare) in a rotating wheel. Beads were washed 5 times for 5 minutes each with WB (10 mM Tris-HCl pH 7.5, 137 mM NaCl, 1 mM EDTA, 1% Triton X-100). Beads were centrifuged at 2,000 x g for 2 minutes after each wash. For elution, samples were boiled for 5 minutes at 95°C in 2x Laemmli buffer.

### Western blot analysis

Cells were lysed in cold RIPA buffer (Cell Signaling Technology), WB5 (8 M urea, 1% SDS) or weak buffer (10 mM PIPES pH 6.8, 100 mM NaCl, 1 mM EGTA, 3 mM MgCl2, 300 mM sucrose, 0.5 mM DTT, 1% Triton X-100) supplemented with 1x protease inhibitor cocktail (Roche), and also in some cases with PhosphoStop 1x (Roche). Lysates were kept on ice for 30 minutes vortexing every 5 minutes and then cleared by centrifugation (25,000 x g, 20 minutes, 4°C). Supernatants were collected and protein content was quantified by BCA protein quantification assay (Pierce). After SDS-PAGE and transfer to nitrocellulose membranes, blocking was performed in 5% milk, or in 5% BSA (Bovine Serum Albumin, Fraction V, Sigma) in PBT (1x PBS, 0.1% Tween-20). In general, primary antibodies were incubated overnight at 4°C and secondary antibodies for 1 hour at room temperature (RT). Antibodies used: anti-LUZP1 (Sigma, 1:1,000), anti-LUZP1 (Proteintech, 1:1,000), anti-CCP110 (Proteintech, 1:1,000), anti-CEP97 (Proteintech, 1:1,000), anti-GFP (Roche, 1:1,000), anti-GAPDH (Proteintech, 1:1,000), anti-FLNA (Merck, 1:1,000), anti-GLI3 (R&D, 1:1,000), HRP-conjugated anti-biotin (Cell Signaling, 1:2,000), anti Myc (Cell Signaling, 1:2,000), anti-actin (Sigma, 1:1,000) and anti-SALL1 (R&D, 1:1,000). Secondary antibodies were anti-goat, anti-mouse or anti-rabbit HRP-conjugates (Jackson Immunoresearch). Proteins were detected using Clarity ECL (BioRad) or Super Signal West Femto (Pierce). Quantification of bands was performed using ImageJ software and normalized against GAPDH or actin levels. At least three independent blots were quantified per experiment.

### Immunostaining

Shh-LIGHT2 cells, hTERT-RPE1, U2OS cells and primary fibroblasts from control and TBS individuals were seeded on 11 mm coverslips (15,000-25,000 cells per well; 24 well-plate). After washing 3 times with cold 1xPBS, cells were fixed with methanol 100% for 10 minutes at −20°C or with 4% PFA supplemented with 0.1% Triton X-100 in PBS for 15 minutes at RT. Then, coverslips were washed 3 times with 1x PBS. Blocking was performed for 1 hour at 37°C in blocking buffer (BB: 2% foetal calf serum, 1% BSA in 1x PBS). Primary antibodies were incubated overnight at 4°C and cells were washed with 1x PBS 3 times. To label the ciliary axoneme and the basal body/pericentriolar region, we used mouse antibodies against acetylated alpha-tubulin (Santa Cruz Biotechnologies, 1:160) and gamma-tubulin (Proteintech, 1:160). Other antibodies include, anti Centrin-2 (CETN2, Biolegend, 1:160), anti-LUZP1 (Sigma, 1:100), anti-LUZP1 (Proteintech, 1:100), anti-CCP110 (Proteintech, 1:200), anti-ODF2 (Atlas, 1:100), anti-PCM-1 (Cell Signaling, 1:100) and anti-Centrobin (Genetex, 1:100).

Donkey anti rat, anti-mouse or anti-rabbit secondary antibodies (Jackson Immunoresearch) conjugated to Alexa 488 or Alexa 568 (1:200), GFP booster (Chromotek, 1:500) and Alexa 568-conjugated phalloidin (Invitrogen 1:500), were incubated for 1 hour at 37°C, followed by nuclear staining with DAPI (10 minutes, 300 ng/ml in PBS; Sigma). Fluorescence imaging was performed using an upright fluorescent microscope (Axioimager D1, Zeiss) or super-resolution microscopy (Leica SP8 Lightning and Zeiss LSM 880 Fast Airyscan) with 63x Plan ApoChromat NA1.4. For cilia measurements and counting, primary cilia from at least fifteen different fluorescent micrographs taken for each experimental condition were analyzed using the ruler tool from Adobe Photoshop. Cilia frequency was obtained dividing the number of total cilia by the number of nuclei on each micrograph. Number of cells per micrograph was similar in both TBS and control fibroblasts. To estimate the level of fluorescence in a determined region, we used the mean intensity obtained by ImageJ. To obtain the signal histograms on Figure 2F, we used the plot profile tool in FIJI.

### qPCR analysis

TBS^275^, control fibroblasts, and Shh-LIGHT2 cells were starved for 24 hours with or without purmorphamine treatment (5 μM; ChemCruz) during 24 hours to induce Shh signaling pathway. Total RNA was obtained with EZNA Total RNA Kit (Omega) and quantified by Nanodrop spectrophotometer. cDNAs were prepared using the SuperScript III First-Strand Synthesis System (Invitrogen) in 10 µl volume per reaction. *LUZP1, GAPDH, Gli1*, *Ptch1*, and *Rplp0* primers were tested for efficiency and products checked for correct size before being used in test samples. qPCR was done using PerfeCTa SYBR Green SuperMix Low (Quantabio). Reactions were performed in 10 µl, adding 1 µl of cDNA and 0.5 µl of each primer (10 µM), in a CFX96 thermocycler (BioRad) using the following protocol: 95°C for 10 minutes and 40 cycles of 95°C for 10 seconds and 55-60°C for 30 seconds. Melting curve analysis was performed for each pair of primers between 65°C and 95°C, with 0.5°C temperature increments every 5 seconds. Relative gene expression data were analyzed using the ΔΔCt method. Reactions were done in triplicates and results were derived from at least three independent experiments normalized to *GAPDH* and *Rplp0* and presented as relative expression levels. Primer sequences: *LUZP1-F:* 5’-GGAATCGGGTAGGAGACACCA-3’; *LUZP1-R:* 5’-TTCCCAGGCAGTTCAGACGGA-3; *GAPDH-F*: 5’-AGCCACATCGCTCAGACAC-3’; *GAPDH-R*: 5’-GCCCAATACGACCAAATCC-3’; *Gli1-F*: 5’-AGCCTTCAGCAATGCCAGTGAC-3’; *Gli1-R*: 5’-

GTCAGGACCATGCACTGTCTTG-3’; *Ptch1-F*: 5’-AAGCCGACTACATGCCAGAG-3’; *Ptch1-R*: 5’-TGATGCCATCTGCGTCTACCAG-3’, *Rplp0-F*: 5’-ACTGGTCTAGGACCCGAGAAG-3’; *Rplp0-R*: 5’-CTCCCACCTTGTCTCCAGTC-3’.

### Luciferase assays

Firefly luciferase expression was measured using the Dual-Luciferase Reporter Assay System (Promega) according to the manufacturer’s instructions. Luminescence was measured and data were normalized to the Renilla luciferase readout. For each construct, luciferase activity upon purmorphamine treatment was divided by the activity of cells before treatment to obtain the fold change value. Experiments were performed with both biological (3x) and technical replicates (n=6).

### Statistical analysis

Statistical analysis was performed using GraphPad 6.0 software. Data were analyzed by Shapiro-Wilk normality test and Levenés test of variance. We used two-tailed unpaired Student’s t-test or Mann Whitney-U tests for comparing two groups, One-way ANOVA or Kruskall-Wallis and the corresponding post-hoc tests for more than two groups and two-way ANOVA to compare more than one variable in more than two groups. P values were represented by asterisks as follows: (*) P-value < 0.05; (**) P-value < 0.01; (***) P-value < 0.001; (****) P-value < 0.0001. Differences were considered significant when P < 0.05.

## DATA AVAILABILITY

The data that support the findings of this study are available from the corresponding author upon reasonable request.

## ACKNOWLEDGEMENTS

We acknowledge A. Cenigaonandia for her assistance in the experiments. We are grateful to the Fundación Inocente, Inocente for their support. We thank the Servicio General de Microscopía Analítica y de Alta Resolución en Biomedicina, SGIker at the UPV/EHU. We also acknowledge funding by grants BFU2017-84653-P (MINECO/FEDER, EU), SEV-2016-0644 (Severo Ochoa Excellence Program), 765445-EU (UbiCODE Program), SAF2017-90900-REDT (UBIRed Program), and IT634-13 (Basque Country Government). Additional support was provided by the Department of Industry, Tourism, and Trade of the Basque Country Government (Elkartek Research Programs) and by the Innovation Technology Department of the Bizkaia County. FE is at Proteomics Platform, member of ProteoRed-ISCIII (PT13/0001/0027) and CIBERehd.

## Contributions

L.B.-B., J.D.S. and R.B designed experiments, analyzed data and wrote the manuscript. L.B.-B., M.G.-S., A.B.-A., C.D., N.M.-M., F.E. and J.D.S. developed experimental protocols, performed experiments, and analyzed data. O.P., R.A. T.C.B., A.Y.T., A.C. and J.A.R. provided scientific resources.

## FIGURE SUPPLEMENT LEGENDS

**Figure 1-figure supplement 1.**
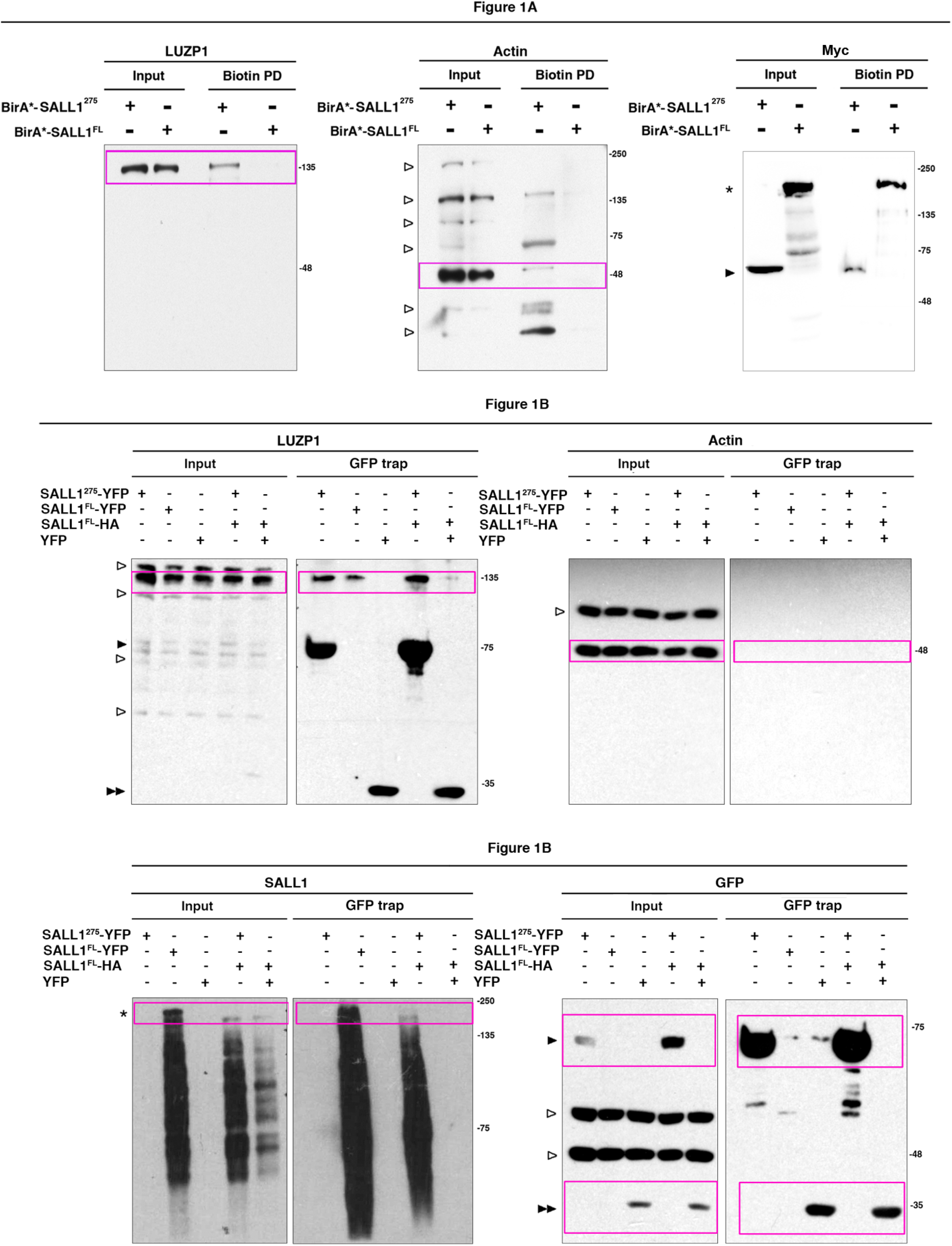
Western blot full pictures for Figure 1. Titles indicate the Figure where each Western blot belongs to; magenta boxes show the region of the gel that was used to build the indicated figures. SALL1^FL^ protein is indicated by one asterisk, SALL1 truncated forms by one black arrowhead, YFP alone by two black arrowheads. Bands from previous probing or unspecific bands are indicated by one empty arrowhead. Molecular weight markers are shown to the right.

**Figure 2-figure supplement 1.**
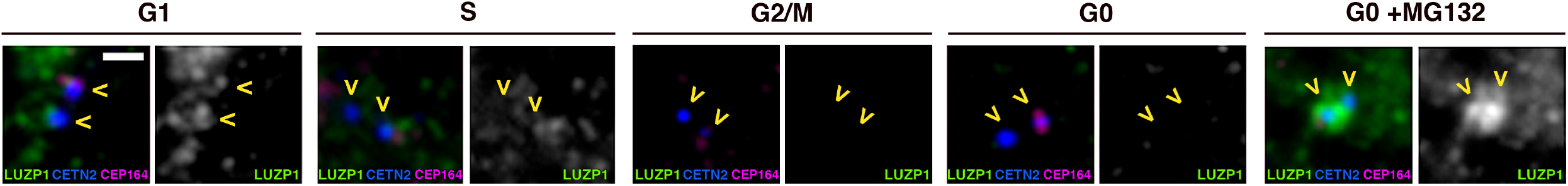
Centrosomal localization of LUZP1 along the cell cycle. Immunofluorescence micrographs showing LUZP1 in the centrosome during cell cycle in RPE1 cells. Cells were treated with mimosine (G1 phase), thymidine (S phase), RO-3306 (G2/M phase) or starved (G0) with and without the proteasome inhibitor MG132. Cells were stained in green with antibodies against endogenous LUZP1, in magenta with CEP164 and in blue with CETN2 (yellow arrowheads). Note a general decrease of LUZP1 during G2/M and upon starvation (G0), which is recovered by MG132 addition. Scale bar 0.5 µm. Images were taken using widefield fluorescence microscopy (Zeiss Axioimager D1, 63x objective).

**Figure 2-Supplementary video 1. LUZP1 localization in the centrosome.** 3D reconstruction of Z-stack micrographs of human control fibroblasts (ESCTRL#2) stained with antibodies against endogenous LUZP1 (green) and acetylated alpha and gamma tubulin to label the cilia and centrosomes, respectively (magenta). Image was taken using Confocal Super-resolution microscopy (LSM 980, Zeiss).

**Figure 3-figure supplement 1.**
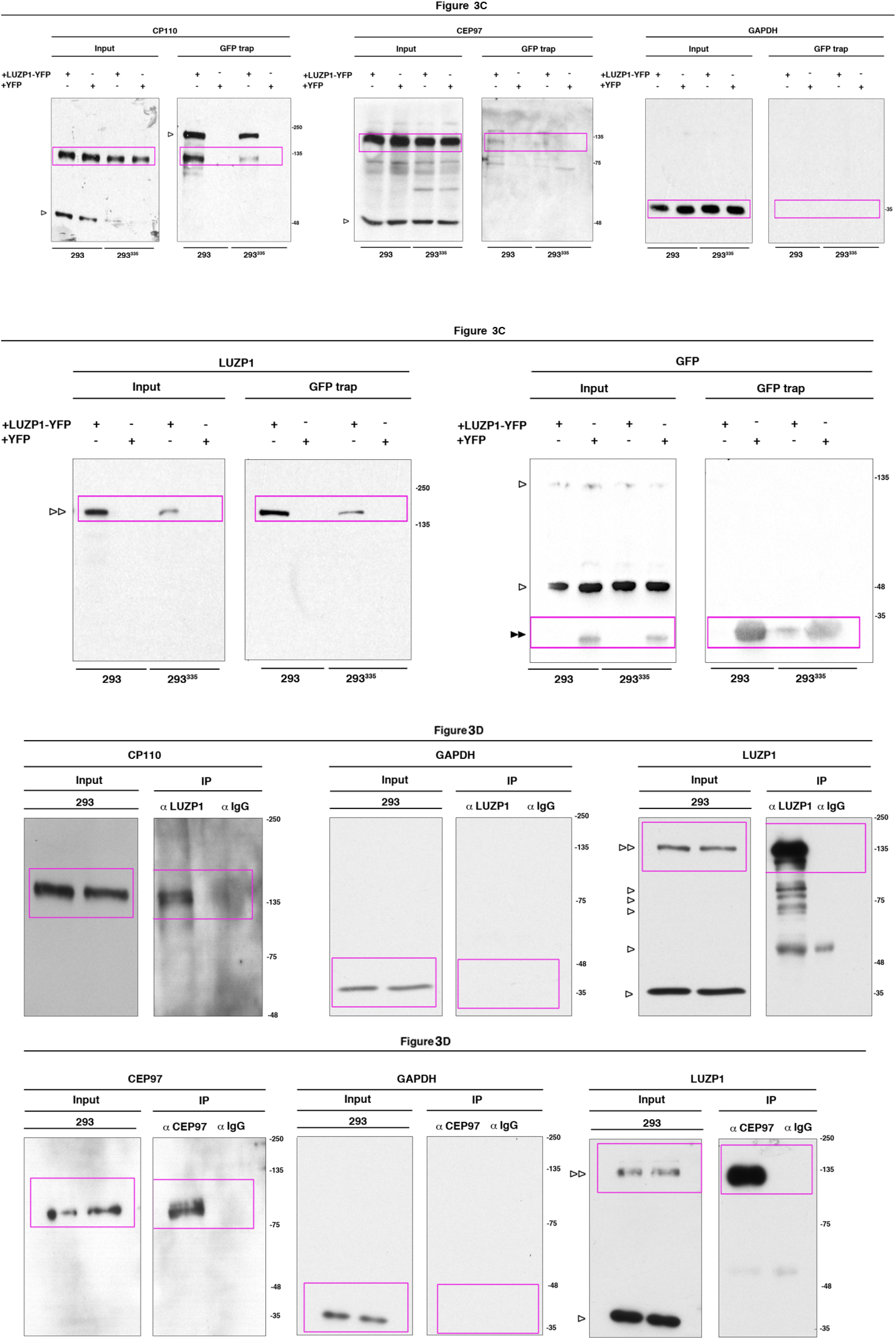
Western blot full pictures for Figure 3. Titles indicate the Figure where each Western blot belongs to; magenta boxes show the region of the gel that was used to build the indicated figures. LUZP1-YFP is indicated by two empty arrowheads and YFP alone by two black arrowheads. Bands from previous probing or unspecific bands are indicated by one empty arrowhead. Molecular weight markers are shown to the right.

**Figure 4-figure supplement 1.**
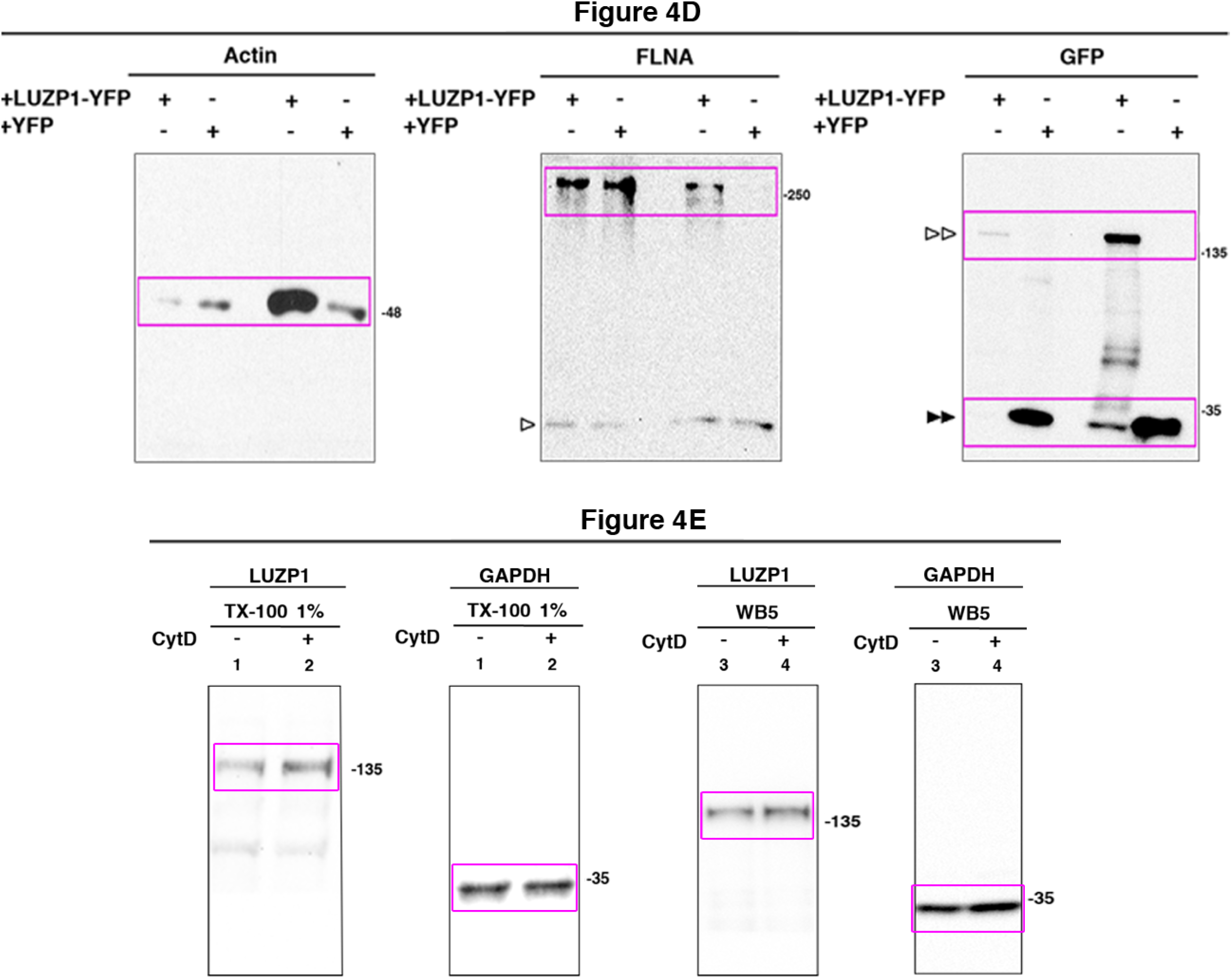
Western blot full pictures for Figure 4. Titles indicate the Figure where each Western blot belongs to; magenta boxes show the region of the gel that was used to build the indicated figures. LUZP1-YFP is indicated by two empty arrowheads and YFP alone by two black arrowheads. Bands from previous probing or unspecific bands are indicated by one empty arrowhead. Molecular weight markers are shown to the right.

**Figure 4-figure supplement 2.**
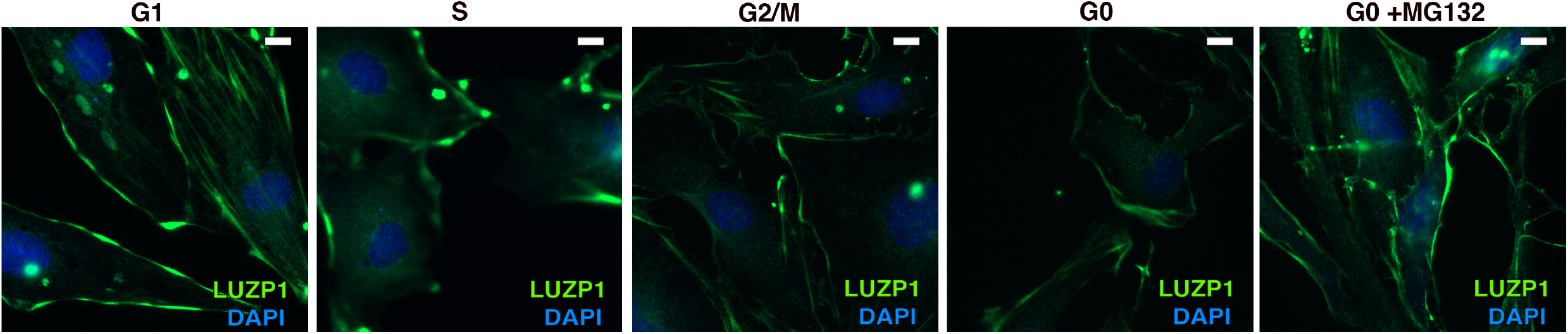
LUZP1 localization at the cytoskeleton along the cell cycle. Immunofluorescence micrographs showing LUZP1 in the whole cell during cell cycle in RPE1 cells. Cells were treated with mimosine (G1 phase), thymidine (S phase), RO-3306 (G2/M phase) or starved (G0) with and without the proteasome inhibitor MG132. Cells were stained in green with antibodies against endogenous LUZP1, and in blue with DAPI. Note a general decrease of LUZP1 during G2/M and upon starvation (G0), which is recovered by MG132 addition. Scale bar 10 µm. Images were taken using widefield fluorescence microscopy (Zeiss Axioimager D1, 63x objective).

**Figure 4-figure supplement 3.**
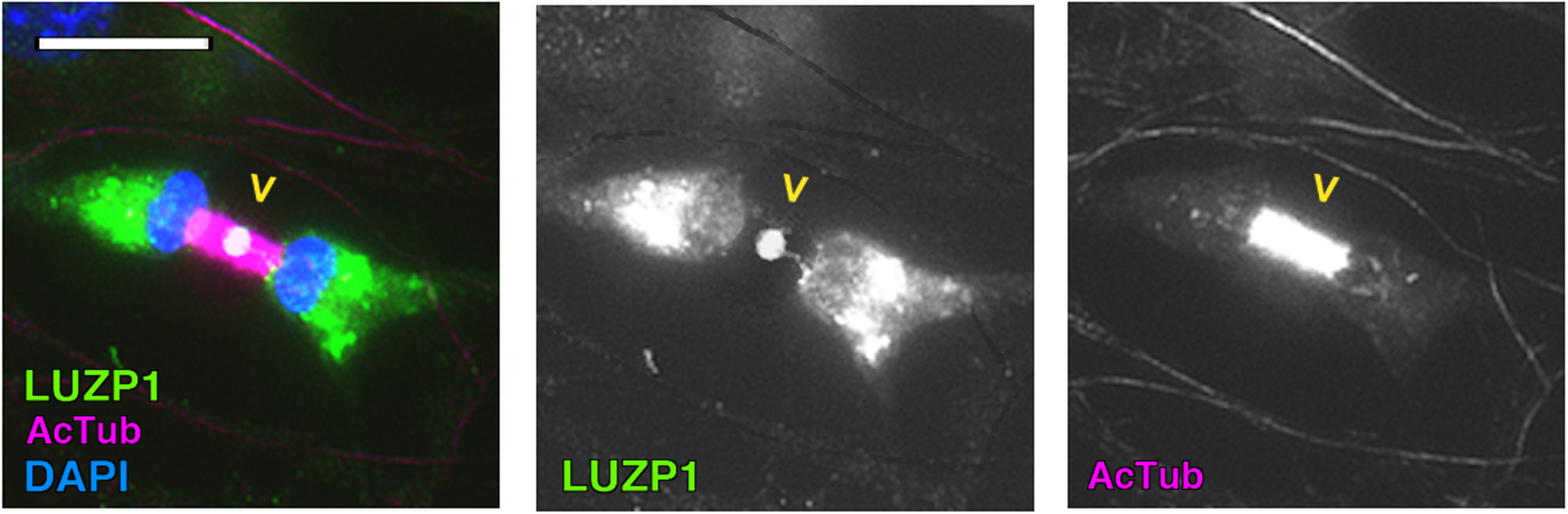
LUZP1 localization at the midbody. Immunofluorescence micrographs of two dividing Shh-LIGHT2 WT cells stained with antibodies against endogenous LUZP1 (green), acetylated tubulin to label microtubules (magenta) and DAPI to label the nuclei (blue). Note the presence of LUZP1 in the midbody (yellow arrowhead). Scale bar 30 µm. Imaging was performed using widefield fluorescence microscopy (Zeiss Axioimager D1, 63x objective).

**Figure 5-figure supplement 1.**
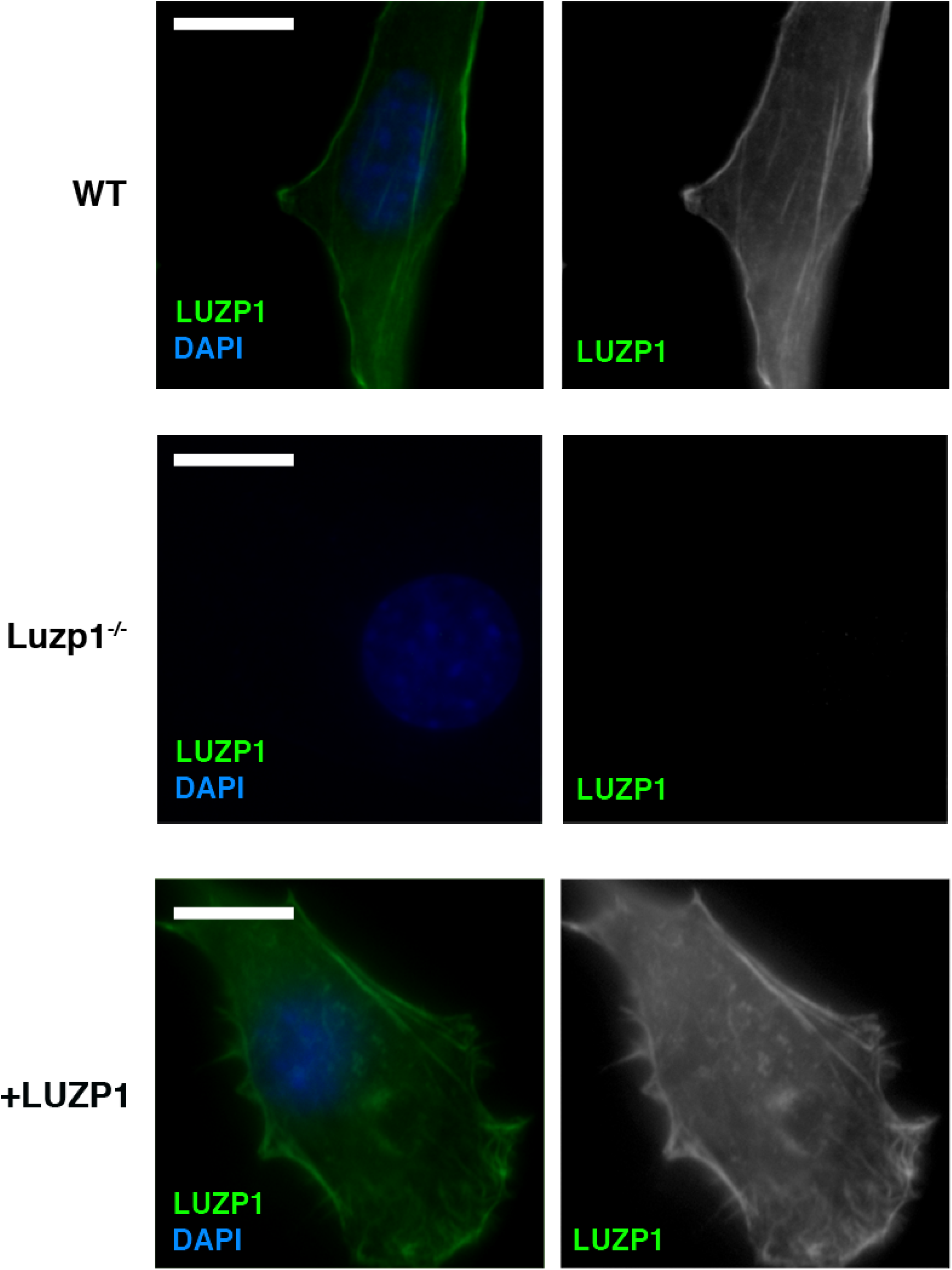
LUZP1 mutant cells and antibody validation at the cytoskeleton. Immunofluorescence micrographs of Shh-LIGHT2 control cells (WT), *Luzp1* depleted Shh-LIGHT2 cells (Luzp1^−/−^) and Luzp1^−/−^ cells rescued with human *LUZP1* (+LUZP1 cells) stained with a specific antibody against endogenous LUZP1 (Sigma, green) and DAPI (blue). Single green channels are shown in black and white. Note the lack of LUZP1 in Luzp1^−/−^ cells. Scale bar, 10 µm.

**Figure 5-figure supplement 2.**
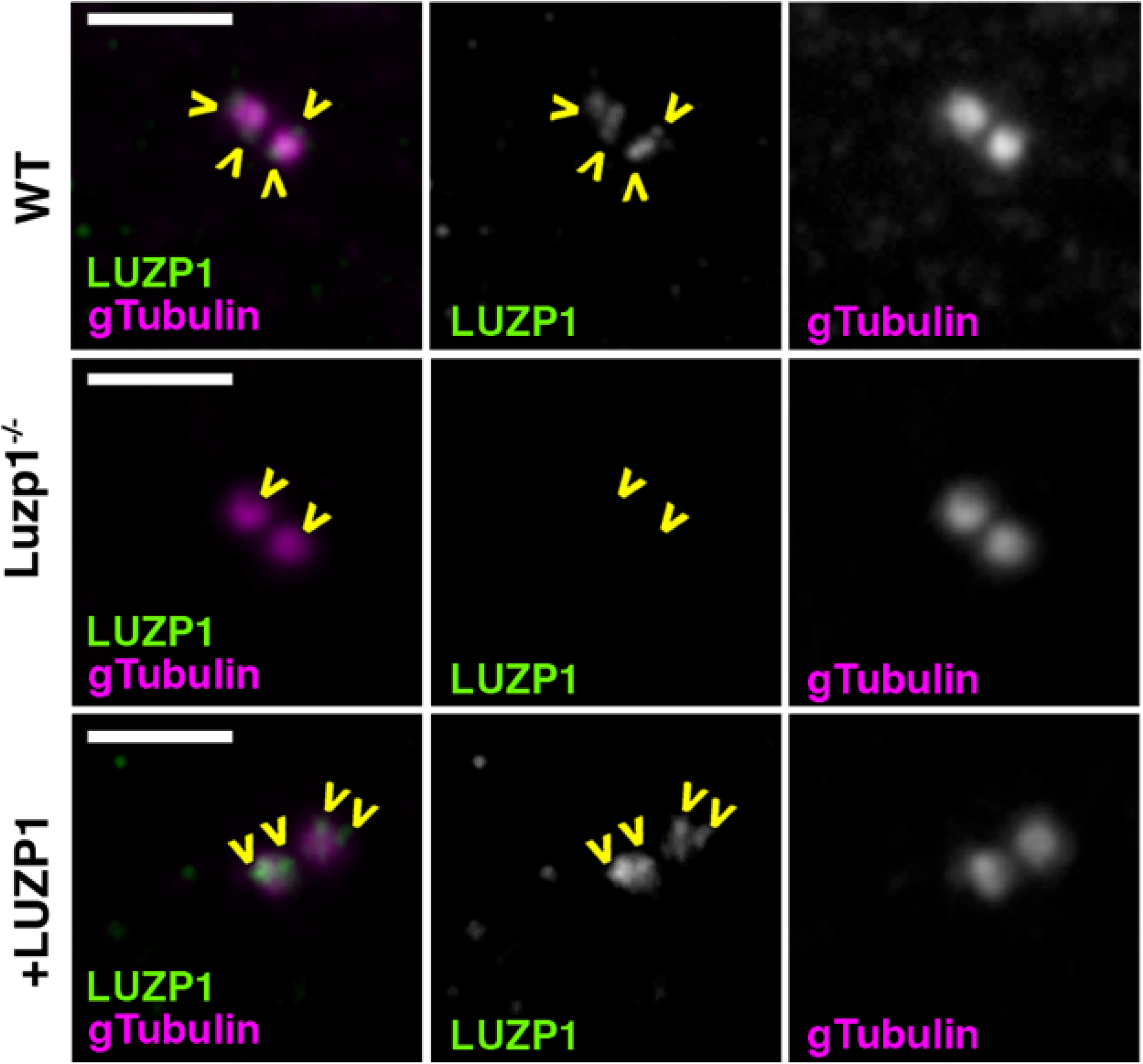
LUZP1 mutant cells and antibody validation at the centrosome. Immunofluorescence micrographs of Shh-LIGHT2 control cells (WT), *Luzp1* depleted Shh-LIGHT2 cells (Luzp1^−/−^) and Luzp1^−/−^ cells stained with antibodies against endogenous LUZP1 (green) and gamma tubulin (magenta). Single green and magenta channels are shown in black and white. Note the lack of LUZP1 in the centrosome in Luzp1^−/−^ cells. Scale bar, 2.5 µm. Images were taken using widefield fluorescence microscopy (Zeiss Axioimager D1, 63x objective).

**Figure 5-figure supplement 2.**
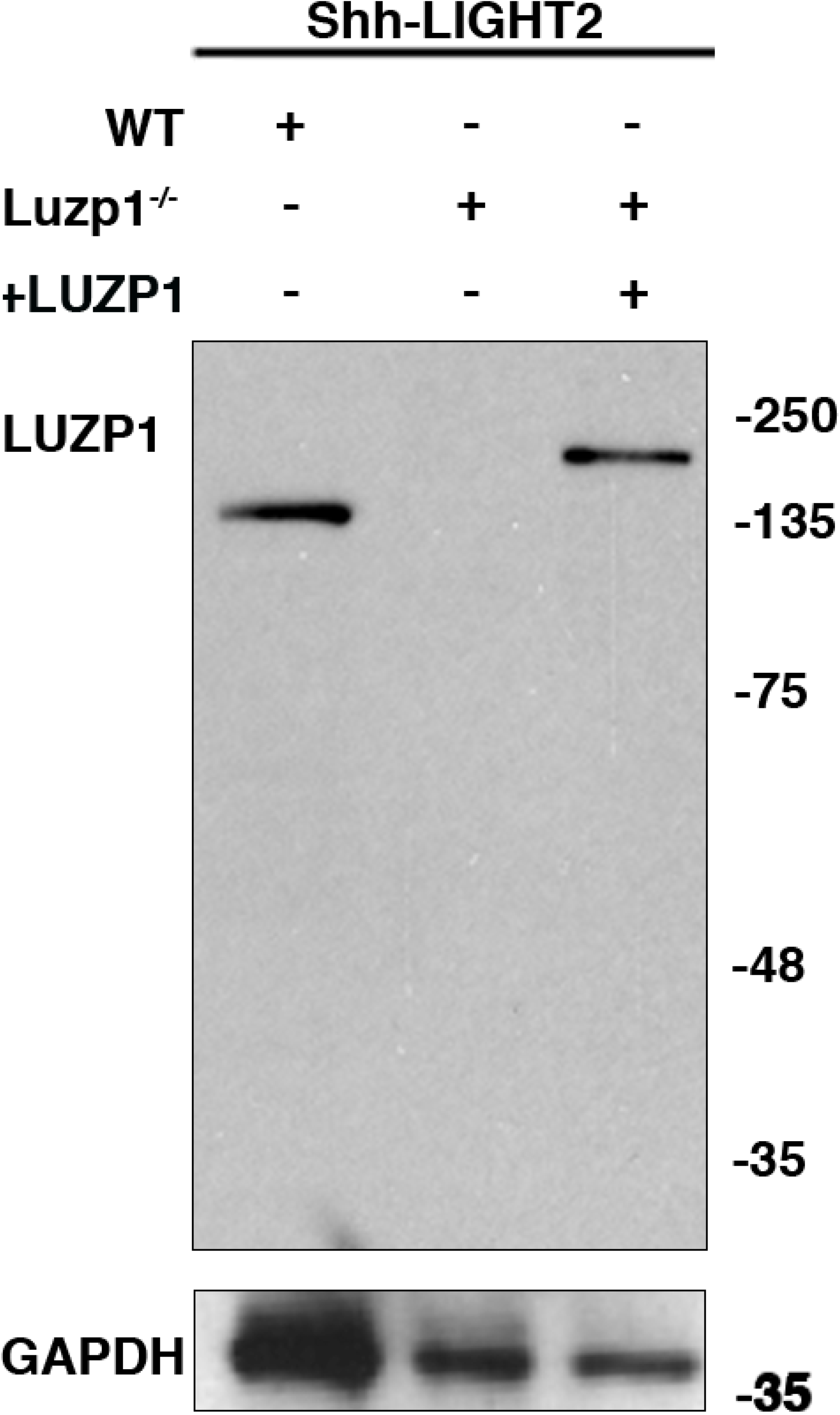
LUZP1 and antibody validation by Western blot. Western blot analysis of total lysates of Shh-LIGHT2 control cells (WT), *Luzp1* depleted Shh-LIGHT2 cells (Luzp1^−/−^) and Luzp1^−/−^ cells using anti-LUZP1 antibodies. Molecular weights in kDa are shown to the right.

**Figure 6-figure supplement 1.**
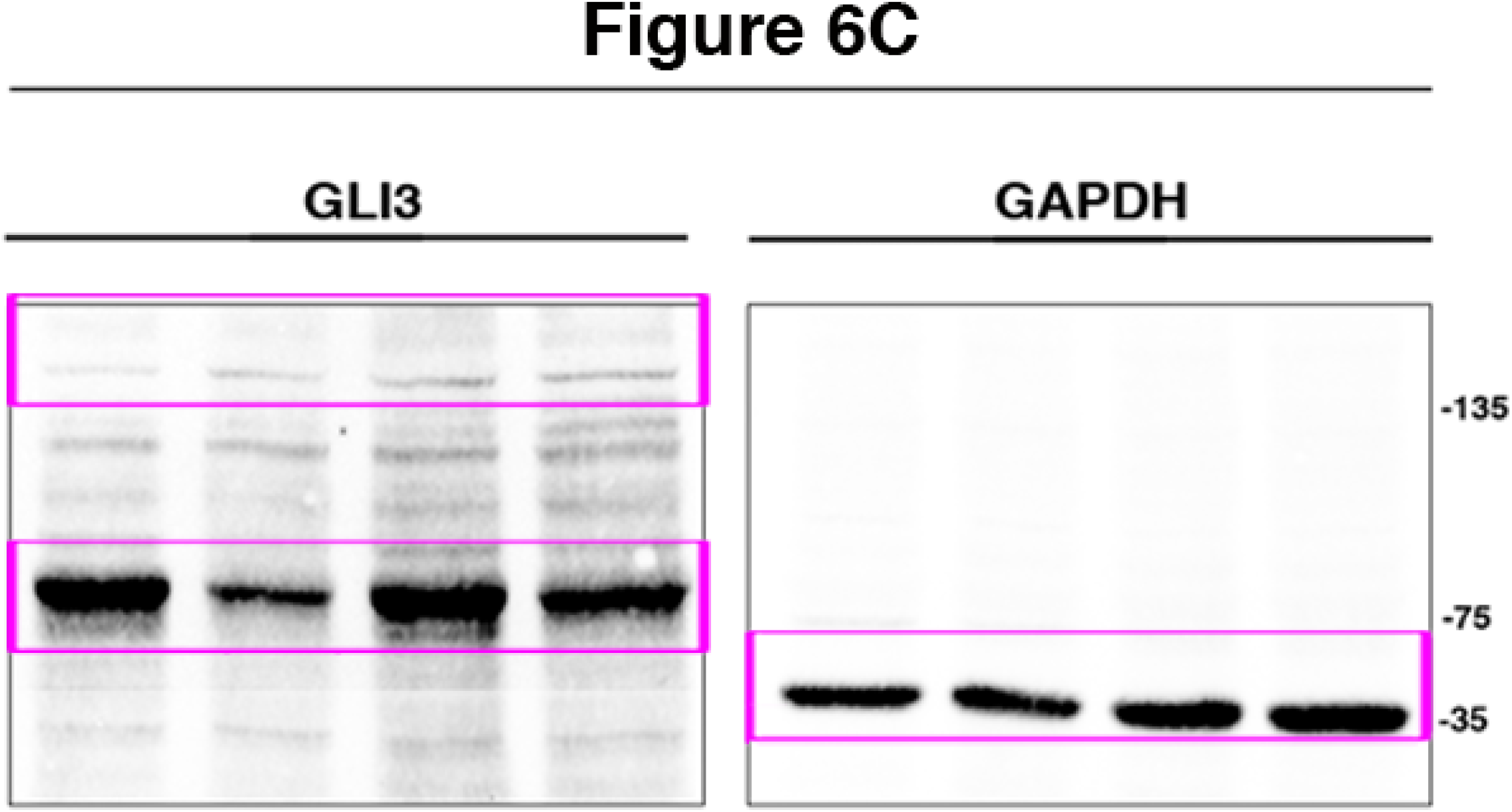
Western blot full pictures for Figure 6. Title indicates the Figure where the Western blot belongs to; magenta boxes show the region of the gel that was used to build the indicated figures. Molecular weight markers are shown to the right.

**Figure 7-figure supplement 1.**
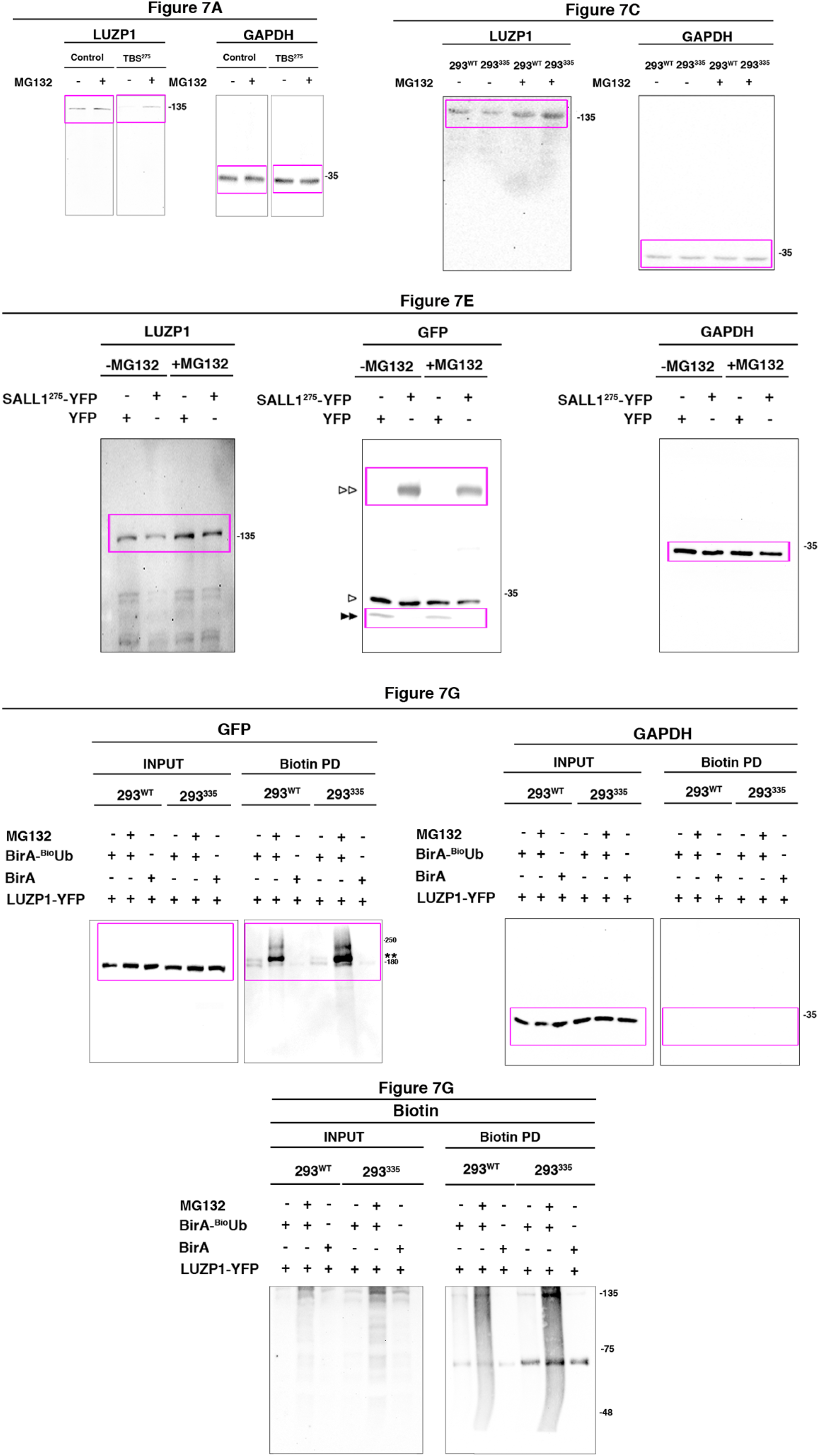
Western blot full pictures for Figure 7. Titles indicate the Figure where each Western blot belongs to; magenta boxes show the region of the gel that was used to build the indicated figures. Ubiquitinated LUZP1 is indicated by two asterisks, LUZP1-YFP by two empty arrowheads and YFP alone by two black arrowheads. Bands from previous probing or unspecific bands are indicated by one empty arrowhead. Molecular weight markers are shown to the right.

**Figure 7-figure supplement 2.**
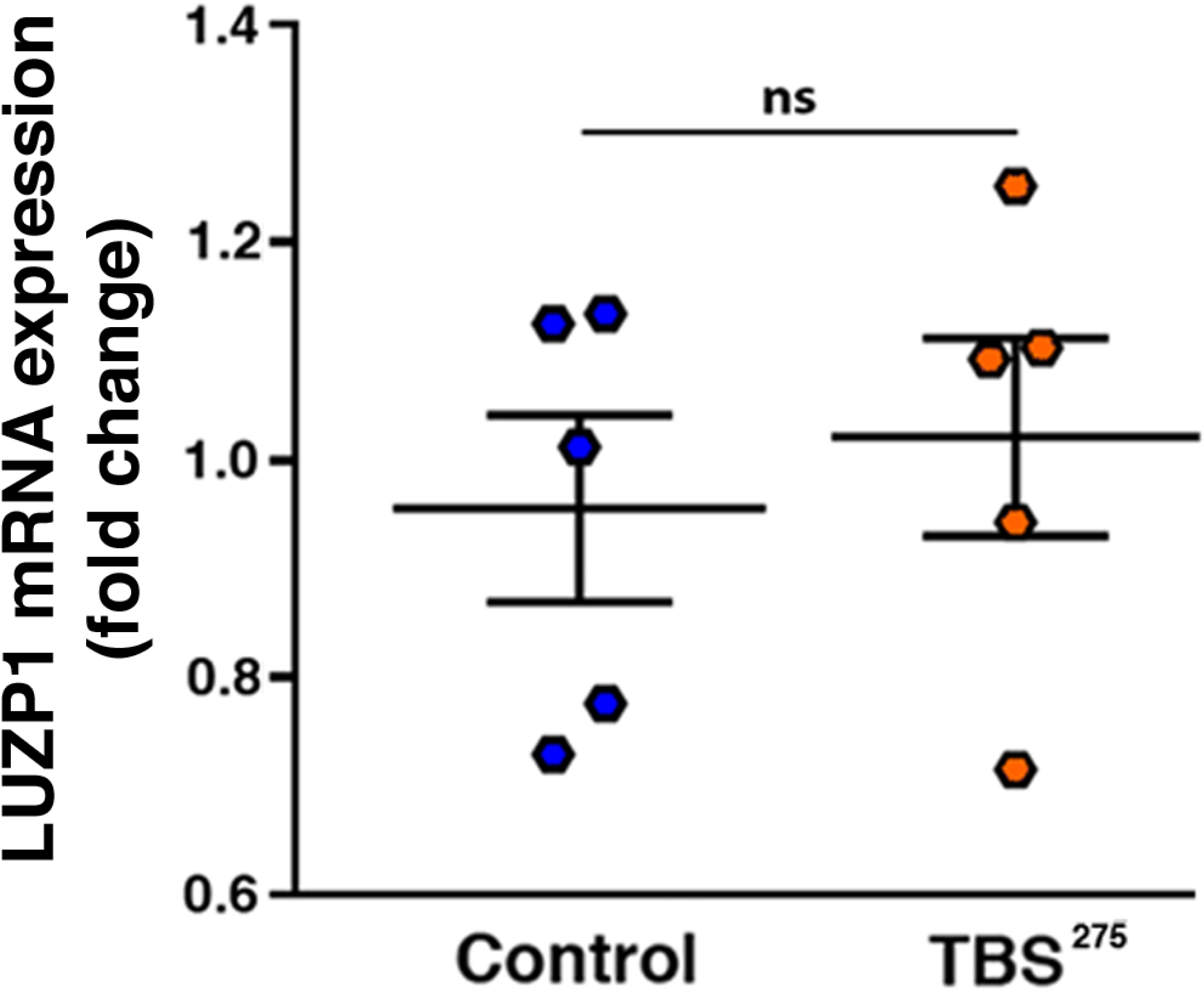
LUZP1 mRNA expression levels. Quantification of *LUZP1* expression in control (ESCTRL2) *vs* TBS^275^ cells by qPCR. Graphs represent Mean and SEM from 5 independent experiments. P-values were calculated using the Mann Whitney test.

## TABLE TITLE AND LEGEND

**Table S1. Identification of LUZP1 interactors by proximity proteomics.**

## REFERENCES

1. Goetz SC, Anderson KV. The primary cilium: a signalling centre during vertebrate development. Nat Rev Genet. 2010;11(5):331–44. 10.1038/nrg2774I.

2. Rezabkova L, Kraatz SH, Akhmanova A, Steinmetz MO, Kammerer RA. Biophysical and Structural Characterization of the Centriolar Protein Cep104 Interaction Network. J Biol Chem. 2016;291(35):18496–504. 10.1074/jbc.M116.739771I.

3. Conduit PT, Wainman A, Raff JW. Centrosome function and assembly in animal cells. Nat Rev Mol Cell Biol. 2015;16(10):611–24. 10.1038/nrm4062I.

4. Vertii A, Hehnly H, Doxsey S. The Centrosome, a Multitalented Renaissance Organelle. Cold Spring Harb Perspect Biol. 2016;8(12). 10.1101/cshperspect.a025049I.

5. Fu J, Glover DM. Structured illumination of the interface between centriole and peri-centriolar material. Open Biol. 2012;2(8):120104. 10.1098/rsob.120104I.

6. Lawo S, Hasegan M, Gupta GD, Pelletier L. Subdiffraction imaging of centrosomes reveals higher-order organizational features of pericentriolar material. Nat Cell Biol. 2012;14(11):1148–58. 10.1038/ncb2591I.

7. Mennella V, Keszthelyi B, McDonald KL, Chhun B, Kan F, Rogers GC, et al. Subdiffraction-resolution fluorescence microscopy reveals a domain of the centrosome critical for pericentriolar material organization. Nat Cell Biol. 2012;14(11):1159–68. 10.1038/ncb2597I.

8. Sonnen KF, Schermelleh L, Leonhardt H, Nigg EA. 3D-structured illumination microscopy provides novel insight into architecture of human centrosomes. Biol Open. 2012;1(10):965–76. 10.1242/bio.20122337I.

9. Kim TS, Zhang L, Il Ahn J, Meng L, Chen Y, Lee E, et al. Molecular architecture of a cylindrical self-assembly at human centrosomes. Nat Commun. 2019;10(1):1151. 10.1038/s41467-019-08838-2I.

10. Gonczy P. Centrosomes and cancer: revisiting a long-standing relationship. Nat Rev Cancer. 2015;15(11):639–52. 10.1038/nrc3995I.

11. Nigg EA, Holland AJ. Once and only once: mechanisms of centriole duplication and their deregulation in disease. Nat Rev Mol Cell Biol. 2018;19(5):297–312. 10.1038/nrm.2017.127I.

12. Spektor A, Tsang WY, Khoo D, Dynlacht BD. Cep97 and CP110 suppress a cilia assembly program. Cell. 2007;130(4):678–90. 10.1016/j.cell.2007.06.027I.

13. Pitaval A, Tseng Q, Bornens M, Thery M. Cell shape and contractility regulate ciliogenesis in cell cycle-arrested cells. J Cell Biol. 2010;191(2):303–12. 10.1083/jcb.201004003I.

14. Fliegauf M, Benzing T, Omran H. When cilia go bad: cilia defects and ciliopathies. Nat Rev Mol Cell Biol. 2007;8(11):880–93. 10.1038/nrm2278I.

15. Hildebrandt F, Benzing T, Katsanis N. Ciliopathies. N Engl J Med. 2011;364(16):1533–43. 10.1056/NEJMra1010172I.

16. Botzenhart EM, Bartalini G, Blair E, Brady AF, Elmslie F, Chong KL, et al. Townes-Brocks syndrome: twenty novel SALL1 mutations in sporadic and familial cases and refinement of the SALL1 hot spot region. Hum Mutat. 2007;28(2):204–5. 10.1002/humu.9476I.

17. Kohlhase J, Wischermann A, Reichenbach H, Froster U, Engel W. Mutations in the SALL1 putative transcription factor gene cause Townes-Brocks syndrome. Nat Genet. 1998;18(1):81–3. 10.1038/ng0198-81I.

18. Bozal-Basterra L, Martin-Ruiz I, Pirone L, Liang Y, Sigurethsson JO, Gonzalez-Santamarta M, et al. Truncated SALL1 Impedes Primary Cilia Function in Townes-Brocks Syndrome. Am J Hum Genet. 2018;102(2):249–65. 10.1016/j.ajhg.2017.12.017I.

19. Wang J, Nakamura F. Identification of Filamin A Mechanobinding Partner II: Fimbacin Is a Novel Actin Cross-Linking and Filamin A Binding Protein. Biochemistry. 2019. 10.1021/acs.biochem.9b00101I.

20. Hein MY, Hubner NC, Poser I, Cox J, Nagaraj N, Toyoda Y, et al. A human interactome in three quantitative dimensions organized by stoichiometries and abundances. Cell. 2015;163(3):712–23. 10.1016/j.cell.2015.09.053I.

21. Nagano T, Morikubo S, Sato M. Filamin A and FILIP (Filamin A-Interacting Protein) regulate cell polarity and motility in neocortical subventricular and intermediate zones during radial migration. J Neurosci. 2004;24(43):9648–57. 10.1523/JNEUROSCI.2363-04.2004I.

22. Gad AK, Nehru V, Ruusala A, Aspenstrom P. RhoD regulates cytoskeletal dynamics via the actin nucleation-promoting factor WASp homologue associated with actin Golgi membranes and microtubules. Mol Biol Cell. 2012;23(24):4807–19. 10.1091/mbc.E12-07-0555I.

23. Hsu CY, Chang NC, Lee MW, Lee KH, Sun DS, Lai C, et al. LUZP deficiency affects neural tube closure during brain development. Biochem Biophys Res Commun. 2008;376(3):466–71. 10.1016/j.bbrc.2008.08.170I.

24. Botzenhart EM, Green A, Ilyina H, Konig R, Lowry RB, Lo IF, et al. SALL1 mutation analysis in Townes-Brocks syndrome: twelve novel mutations and expansion of the phenotype. Hum Mutat. 2005;26(3):282. 10.1002/humu.9362I.

25. Surka WS, Kohlhase J, Neunert CE, Schneider DS, Proud VK. Unique family with Townes-Brocks syndrome, SALL1 mutation, and cardiac defects. Am J Med Genet. 2001;102(3):250–7.

26. Klena N, Gabriel G, Liu X, Yagi H, Li Y, Chen Y, et al. Role of Cilia and Left-Right Patterning in Congenital Heart Disease. In: Nakanishi T, Markwald RR, Baldwin HS, Keller BB, Srivastava D, Yamagishi H, editors. Etiology and Morphogenesis of Congenital Heart Disease: From Gene Function and Cellular Interaction to Morphology. Tokyo 2016. p. 67–79.

27. Toomer KA, Yu M, Fulmer D, Guo L, Moore KS, Moore R, et al. Primary cilia defects causing mitral valve prolapse. Sci Transl Med. 2019;11(493). 10.1126/scitranslmed.aax0290I.

28. Lee MW, Chang AC, Sun DS, Hsu CY, Chang NC. Restricted expression of LUZP in neural lineage cells: a study in embryonic stem cells. J Biomed Sci. 2001;8(6):504–11. 10.1159/000046172I.

29. Sun DS, Chang AC, Jenkins NA, Gilbert DJ, Copeland NG, Chang NC. Identification, molecular characterization, and chromosomal localization of the cDNA encoding a novel leucine zipper motif-containing protein. Genomics. 1996;36(1):54–62. 10.1006/geno.1996.0425I.

30. Campbell K. Dorsal-ventral patterning in the mammalian telencephalon. Curr Opin Neurobiol. 2003;13(1):50–6.

31. Copp AJ. Neurulation in the cranial region--normal and abnormal. J Anat. 2005;207(5):623–35. 10.1111/j.1469-7580.2005.00476.xI.

32. Fuccillo M, Joyner AL, Fishell G. Morphogen to mitogen: the multiple roles of hedgehog signalling in vertebrate neural development. Nat Rev Neurosci. 2006;7(10):772–83. 10.1038/nrn1990I.

33. Branon TC, Bosch JA, Sanchez AD, Udeshi ND, Svinkina T, Carr SA, et al. Efficient proximity labeling in living cells and organisms with TurboID. Nat Biotechnol. 2018;36(9):880–7. 10.1038/nbt.4201I.

34. Alves-Cruzeiro JM, Nogales-Cadenas R, Pascual-Montano AD. CentrosomeDB: a new generation of the centrosomal proteins database for Human and Drosophila melanogaster. Nucleic Acids Res. 2014;42(Database issue):D430–6. 10.1093/nar/gkt1126I.

35. Gupta GD, Coyaud E, Goncalves J, Mojarad BA, Liu Y, Wu Q, et al. A Dynamic Protein Interaction Landscape of the Human Centrosome-Cilium Interface. Cell. 2015;163(6):1484–99. 10.1016/j.cell.2015.10.065I.

36. Uhlen M, Fagerberg L, Hallstrom BM, Lindskog C, Oksvold P, Mardinoglu A, et al. Proteomics. Tissue-based map of the human proteome. Science. 2015;347(6220):1260419. 10.1126/science.1260419I.

37. Taipale J, Chen JK, Cooper MK, Wang B, Mann RK, Milenkovic L, et al. Effects of oncogenic mutations in Smoothened and Patched can be reversed by cyclopamine. Nature. 2000;406(6799):1005–9. 10.1038/35023008I.

38. Goetz SC, Liem KF, Jr., Anderson KV. The spinocerebellar ataxia-associated gene Tau tubulin kinase 2 controls the initiation of ciliogenesis. Cell. 2012;151(4):847–58. 10.1016/j.cell.2012.10.010I.

39. Kleylein-Sohn J, Westendorf J, Le Clech M, Habedanck R, Stierhof YD, Nigg EA. Plk4-induced centriole biogenesis in human cells. Dev Cell. 2007;13(2):190–202. 10.1016/j.devcel.2007.07.002I.

40. Prosser SL, Morrison CG. Centrin2 regulates CP110 removal in primary cilium formation. J Cell Biol. 2015;208(6):693–701. 10.1083/jcb.201411070I.

41. Tsang WY, Bossard C, Khanna H, Peranen J, Swaroop A, Malhotra V, et al. CP110 suppresses primary cilia formation through its interaction with CEP290, a protein deficient in human ciliary disease. Dev Cell. 2008;15(2):187–97. 10.1016/j.devcel.2008.07.004I.

42. Huangfu D, Liu A, Rakeman AS, Murcia NS, Niswander L, Anderson KV. Hedgehog signalling in the mouse requires intraflagellar transport proteins. Nature. 2003;426(6962):83–7. 10.1038/nature02061I.

43. Yin Y, Bangs F, Paton IR, Prescott A, James J, Davey MG, et al. The Talpid3 gene (KIAA0586) encodes a centrosomal protein that is essential for primary cilia formation. Development. 2009;136(4):655–64. 10.1242/dev.028464I.

44. Pirone L, Xolalpa W, Sigurethsson JO, Ramirez J, Perez C, Gonzalez M, et al. A comprehensive platform for the analysis of ubiquitin-like protein modifications using in vivo biotinylation. Sci Rep. 2017;7:40756. 10.1038/srep40756I.

45. Gheiratmand L, Coyaud E, Gupta GD, Laurent EM, Hasegan M, Prosser SL, et al. Spatial and proteomic profiling reveals centrosome-independent features of centriolar satellites. EMBO J. 2019. 10.15252/embj.2018101109I.

46. Bernabe-Rubio M, Andres G, Casares-Arias J, Fernandez-Barrera J, Rangel L, Reglero-Real N, et al. Novel role for the midbody in primary ciliogenesis by polarized epithelial cells. J Cell Biol. 2016;214(3):259–73. 10.1083/jcb.201601020I.

47. Akimov V, Barrio-Hernandez I, Hansen SVF, Hallenborg P, Pedersen AK, Bekker-Jensen DB, et al. UbiSite approach for comprehensive mapping of lysine and N-terminal ubiquitination sites. Nat Struct Mol Biol. 2018;25(7):631–40. 10.1038/s41594-018-0084-yI.

48. Mertins P, Qiao JW, Patel J, Udeshi ND, Clauser KR, Mani DR, et al. Integrated proteomic analysis of post-translational modifications by serial enrichment. Nat Methods. 2013;10(7):634–7. 10.1038/nmeth.2518I.

49. Povlsen LK, Beli P, Wagner SA, Poulsen SL, Sylvestersen KB, Poulsen JW, et al. Systems-wide analysis of ubiquitylation dynamics reveals a key role for PAF15 ubiquitylation in DNA-damage bypass. Nat Cell Biol. 2012;14(10):1089–98. 10.1038/ncb2579I.

50. Udeshi ND, Svinkina T, Mertins P, Kuhn E, Mani DR, Qiao JW, et al. Refined preparation and use of anti-diglycine remnant (K-epsilon-GG) antibody enables routine quantification of 10,000s of ubiquitination sites in single proteomics experiments. Mol Cell Proteomics. 2013;12(3):825–31. 10.1074/mcp.O112.027094I.

51. Wagner SA, Beli P, Weinert BT, Scholz C, Kelstrup CD, Young C, et al. Proteomic analyses reveal divergent ubiquitylation site patterns in murine tissues. Mol Cell Proteomics. 2012;11(12):1578–85. 10.1074/mcp.M112.017905I.

52. D’Angiolella V, Donato V, Vijayakumar S, Saraf A, Florens L, Washburn MP, et al. SCF(Cyclin F) controls centrosome homeostasis and mitotic fidelity through CP110 degradation. Nature. 2010;466(7302):138–42. 10.1038/nature09140I.

53. Hossain D, Javadi Esfehani Y, Das A, Tsang WY. Cep78 controls centrosome homeostasis by inhibiting EDD-DYRK2-DDB1(Vpr)(BP). EMBO Rep. 2017;18(4):632–44. 10.15252/embr.201642377I.

54. Li J, D’Angiolella V, Seeley ES, Kim S, Kobayashi T, Fu W, et al. USP33 regulates centrosome biogenesis via deubiquitination of the centriolar protein CP110. Nature. 2013;495(7440):255–9. 10.1038/nature11941I.

55. Boisvieux-Ulrich E, Laine MC, Sandoz D. Cytochalasin D inhibits basal body migration and ciliary elongation in quail oviduct epithelium. Cell Tissue Res. 1990;259(3):443–54.

56. Euteneuer U, Schliwa M. Evidence for an involvement of actin in the positioning and motility of centrosomes. J Cell Biol. 1985;101(1):96–103.

57. Kim J, Lee JE, Heynen-Genel S, Suyama E, Ono K, Lee K, et al. Functional genomic screen for modulators of ciliogenesis and cilium length. Nature. 2010;464(7291):1048–51. 10.1038/nature08895I.

58. Kim J, Jo H, Hong H, Kim MH, Kim JM, Lee JK, et al. Actin remodelling factors control ciliogenesis by regulating YAP/TAZ activity and vesicle trafficking. Nat Commun. 2015;6:6781. 10.1038/ncomms7781I.

59. Kang GM, Han YM, Ko HW, Kim J, Oh BC, Kwon I, et al. Leptin Elongates Hypothalamic Neuronal Cilia via Transcriptional Regulation and Actin Destabilization. J Biol Chem. 2015;290(29):18146–55. 10.1074/jbc.M115.639468I.

60. Hernandez-Hernandez V, Pravincumar P, Diaz-Font A, May-Simera H, Jenkins D, Knight M, et al. Bardet-Biedl syndrome proteins control the cilia length through regulation of actin polymerization. Hum Mol Genet. 2013;22(19):3858–68. 10.1093/hmg/ddt241I.

61. Drummond ML, Li M, Tarapore E, Nguyen TTL, Barouni BJ, Cruz S, et al. Actin polymerization controls cilia-mediated signaling. J Cell Biol. 2018;217(9):3255–66. 10.1083/jcb.201703196I.

62. Cao J, Shen Y, Zhu L, Xu Y, Zhou Y, Wu Z, et al. miR-129-3p controls cilia assembly by regulating CP110 and actin dynamics. Nat Cell Biol. 2012;14(7):697–706. 10.1038/ncb2512I.

63. Phua SC, Chiba S, Suzuki M, Su E, Roberson EC, Pusapati GV, et al. Dynamic Remodeling of Membrane Composition Drives Cell Cycle through Primary Cilia Excision. Cell. 2017;168(1-2):264–79 e15. 10.1016/j.cell.2016.12.032I.

64. Nager AR, Goldstein JS, Herranz-Perez V, Portran D, Ye F, Garcia-Verdugo JM, et al. An Actin Network Dispatches Ciliary GPCRs into Extracellular Vesicles to Modulate Signaling. Cell. 2017;168(1-2):252–63 e14. 10.1016/j.cell.2016.11.036I.

65. Izawa I, Goto H, Kasahara K, Inagaki M. Current topics of functional links between primary cilia and cell cycle. Cilia. 2015;4:12. 10.1186/s13630-015-0021-1I.

66. Wallingford JB. Neural tube closure and neural tube defects: studies in animal models reveal known knowns and known unknowns. Am J Med Genet C Semin Med Genet. 2005;135C(1):59–68. 10.1002/ajmg.c.30054I.

67. Sadler TW, Greenberg D, Coughlin P, Lessard JL. Actin distribution patterns in the mouse neural tube during neurulation. Science. 1982;215(4529):172–4.

68. Murdoch JN, Copp AJ. The relationship between sonic Hedgehog signaling, cilia, and neural tube defects. Birth Defects Res A Clin Mol Teratol. 2010;88(8):633–52. 10.1002/bdra.20686I.

69. Kohlhase J. Townes-Brocks Syndrome. In: Pagon RA, Adam MP, Ardinger HH, Wallace SE, Amemiya A, Bean LJH, et al., editors. GeneReviews. Seattle (WA) 2007.

70. Bohm J, Buck A, Borozdin W, Mannan AU, Matysiak-Scholze U, Adham I, et al. Sall1, sall2, and sall4 are required for neural tube closure in mice. Am J Pathol. 2008;173(5):1455–63. 10.2353/ajpath.2008.071039I.

71. Roux KJ, Kim DI, Raida M, Burke B. A promiscuous biotin ligase fusion protein identifies proximal and interacting proteins in mammalian cells. J Cell Biol. 2012;196(6):801–10. 10.1083/jcb.201112098I.

72. Meier F, Beck S, Grassl N, Lubeck M, Park MA, Raether O, et al. Parallel Accumulation-Serial Fragmentation (PASEF): Multiplying Sequencing Speed and Sensitivity by Synchronized Scans in a Trapped Ion Mobility Device. J Proteome Res. 2015;14(12):5378–87. 10.1021/acs.jproteome.5b00932I.

73. Reimand J, Arak T, Adler P, Kolberg L, Reisberg S, Peterson H, et al. g:Profiler-a web server for functional interpretation of gene lists (2016 update). Nucleic Acids Res. 2016;44(W1):W83–9. 10.1093/nar/gkw199I.

